# InsectDCT: A generalized pipeline for detection, taxonomic classification, and tracking of insects in camera-trap recordings

**DOI:** 10.64898/2026.07.07.736939

**Authors:** Kim Bjerge, Simon F. A. Wogram, Pau Enric Serra-Marin, Otar Sakhiashvili, Andreas Bonnerup Jepsen, Toke T. Høye

## Abstract

Automated monitoring of insect pollinators in natural environments with insect camera traps and trained deep learning algorithms can provide large-scale, temporally resolved observations for insect ecological studies. However, efficient and accurate recognition of insects in recorded images or videos is challenging, particularly when small insects appear against complex backgrounds with diverse vegetation communities.

In this work, we present *InsectDCT*, an AI-based pipeline for automated detection, hierarchical classification, and tracking of insects during daylight in footage of natural vegetation, tested in different environments. The *InsectDCT* pipeline consists of three levels: insect Detection and localization, hierarchical taxonomic Classification, and spatio-temporal Tracking. In the first stage, insects are detected in time-lapse images or video recordings using the You Only Look Once (YOLO11) object detection architecture. Detection performance is improved using motion-enhanced images, which improve robustness in cluttered environments. The detector is trained on an extensive dataset of more than 60,000 images, and the classifier is trained on 90,358 insect images collected using camera traps deployed across a wide range of plant families and floral habitats. It achieves the best F1-score of 0.88 with motion-enhanced images. In the second stage, detected insects are classified using a hierarchical taxonomy-aware classification framework that covers 80 taxonomic groups. Classification is performed at multiple taxonomic ranks, including order, family, and genus/species, achieving F1-scores of 0.90, 0.84, and 0.76 at each respective rank. It allows for coarse and fine-grained ecological analyzes while accounting for varying levels of visual ambiguity. In the third stage, a multi-object tracking module is applied to high temporal-resolution image sequences and video data to associate detections across consecutive frames and generate track-level observations. It achieves a MOTA of 76.1% and IDF1 of 74.2%. *InsectDCT* code and all datasets are made publicly available.

## 1. Introduction

Insects are the most diverse animal group (Zhang, 2011; Stork, 2018), contributing to pollination, herbivory, decomposition, biological control, and food-web dynamics (Losey and Vaughan, 2006; Potts et al., 2010; Cardoso et al., 2020). At the same time, mounting evidence indicates substantial declines in many insect taxa and regions, creating an urgent need for monitoring systems that can resolve changes in abundance, phenology, and community composition at ecologically significant spatial and temporal scales (Potts et al., 2010; Wagner et al., 2021). Conventional approaches, including direct field observations, trapping, and expert-based taxonomic identification, remain indispensable, but are labor-intensive, spatially constrained, often destructive, and difficult to scale across many sites and seasons (Montgomery et al., 2021; Didham et al., 2020). These limitations restrict our ability to continuously monitor insect populations and link observed changes with land use, climate, habitat management, and insect-plant interactions (Didham et al., 2020; Montgomery et al., 2021; Cardoso et al., 2020; van Klink et al., 2022).

The ongoing global decline in insect populations has coincided with rapid advances in big data, machine learning, and low-cost portable technologies, collectively driving the development of automated monitoring pipelines for diurnal (Jain et al., 2025; Sittinger et al., 2024; Smith et al., 2026) and nocturnal (Bjerge et al., 2021b; Alison et al., 2022; Geissmann et al., 2022) insects. Camera-based systems are particularly promising because they can record insects non-lethally, repeatedly, and under field conditions, generating highly taxonomically resolved information on abundance, activity patterns, phenology, and diversity of insects that would be difficult to obtain manually (Barlow and O’Neill, 2020; Høye et al., 2021; Besson et al., 2022; van Klink et al., 2022; Gillespie et al., 2025). Current approaches vary widely in their ecological context: some monitor the abundance of insects using artificial substrates such as colored panels or sticky surfaces (Sittinger et al., 2024; Geissmann et al., 2022), while others detect insects against natural backgrounds, such as flowers, leaves, or fruits (Droissart et al., 2021; Bjerge et al., 2023a, 2024; Roosjen et al., 2020; Molina-Rotger et al., 2023). These systems have already accelerated the collection of data on ecological interactions, including pollination and herbivory (Alison et al., 2022; Serra-Marin et al., 2025; Wang et al., 2024).

Despite this progress, most existing pipelines remain tailored to specific study systems. Models are commonly trained on a limited set of camera types, plant backgrounds, recording distances, or target taxa, and their performance typically drops when applied to new geographic regions, plant assemblages, image characteristics, or insect communities (Bjerge et al., 2023c, 2024; Smith et al., 2026). This lack of transferability creates a practical bottleneck: Users who want to deploy automated monitoring in a new system often need to collect, annotate, and train on a local dataset before robust inference is possible. This repeated development of site-specific models slows adoption, duplicates effort between projects, and limits comparability among studies. Therefore, a more generalizable framework, trained on a diverse set of backgrounds, camera systems, and pollinator communities is needed to make automated insect monitoring more reusable in ecological contexts (Beery et al., 2018; Høye et al., 2021; Svenning et al., 2026).

A second limitation concerns taxonomic resolution. For many questions in conservation, pollination ecology, and community ecology, detecting an “insect” or assigning a broad order is insufficient. Ecologically meaningful inference often depends on the distinction of families, genera, or species, especially for focal pollinator groups such as bees, hoverflies, and butterflies (Høye et al., 2021; van Klink et al., 2022; Serra-Marin et al., 2025). Fine-grained insect classification in field imagery is difficult because organisms are often small, partially occluded, blurred, variably oriented, or visually similar (Høye et al., 2021; Bjerge et al., 2023c; Jain et al., 2025). Moreover, training datasets typically have long-tailed class imbalance, with many taxa represented by only a few images (Bjerge et al., 2023c). Hierarchical classification provides a biologically natural solution by allowing predictions to remain useful at higher taxonomic ranks when species-level classification is uncertain, but hierarchy-aware classification is still uncommon in operational insect-monitoring pipelines (Bjerge et al., 2023c).

A third challenge is the detection of small objects in complex natural scenes. Camera-trap recognition has long been affected by poor lighting, occlusion, camouflage, blur, and domain shifts between locations (Beery et al., 2018; Oliver et al., 2023). Insect camera traps intensify these problems because target organisms may occupy only a few dozen pixels, overlap with visually similar floral structures, or move out of focus in three dimensional vegetation (Bjerge et al., 2023b, 2024; Ratnayake et al., 2021). Identification can be improved by placing cameras very close to flowers, restricting the field of view, or accepting only relatively large detections (Preti et al., 2021; Ratnayake et al., 2021; Smith et al., 2026). Although these design choices can improve the accuracy of the classification, they will reduce the taxonomic coverage of small insects (Preti et al., 2021; Ratnayake et al., 2021; Smith et al., 2026; Varga-Szilay et al., 2024). A restricted field of view can also affect tracking, because insects are observed for shorter periods, leave the image frame more quickly, and generate fewer consecutive detections from which individual trajectories and behavioral information can be reconstructed (Ratnayake et al., 2021; Bjerge et al., 2021a, 2025). A detector optimized for small insects is therefore essential for monitoring smaller taxa and for recording wider scenes without losing sensitivity, while also supporting more reliable tracking across longer image sequences.

Here we present *InsectDCT*, a generalized pipeline for insect detection, hierarchical taxonomic classification, and tracking in camera recordings from natural and semi-natural floral environments. The framework addresses three linked problems in automated insect monitoring: poor transferability across new field conditions, limited fine-grained taxonomic resolution, and weak performance on small or distant insects. The framework integrates motion-informed object detection, hierarchy-based classification with anomaly handling, and track-by-detection to produce taxonomically annotated insect observations from time-lapse and video recordings (Bjerge et al., 2023b,c, 2025). We trained models on various datasets that contain many floral backgrounds, camera systems, and insect taxa. This design of a general-purpose framework aims to reduce the need for each research group to train a bespoke algorithm from scratch, while still allowing future extension to additional taxa from different geographical regions. In addition, we provide a large public training and evaluation resource for insect detection and classification in natural camera-trap imagery. By releasing both the code and the underlying datasets, *InsectDCT* can serve as a ready-to-use monitoring pipeline and also as a reference and training resource for future algorithms. By combining heterogeneous training data, small-object-oriented detection, hierarchical taxonomic classification, tracking, and open resources, *InsectDCT* provides a reusable computational foundation for scalable insect monitoring in ecological and conservation research.

### 1.1. Contribution

In this work, we propose a flexible and efficient processing pipeline for analyzing image recordings from insect camera traps in natural and semi-natural environments, enabling the detection, classification of insects at multiple taxonomic levels (including order, family, and genus/species) and tracking. A comprehensive dataset is presented, comprising images collected for training detection and classification models across a wide range of insects, flowering plants, and heterogeneity of backgrounds.

We evaluate the performance of the proposed pipeline in terms of precision, recall, and computational efficiency, and assess its processing speed on two edge computing devices: the Raspberry Pi 4 and Raspberry Pi 5. The proposed pipeline supports both time-lapse camera recording strategies and video-based tracking, enabling generic and robust monitoring of insects in natural environments.

In summary, our objectives for this work are the following:

- a generalized insect detection model trained on highly heterogeneous natural backgrounds.
- a hierarchical taxonomic classification model with anomaly-detection.
- to create an integrated tracking framework for insect monitoring in time-lapse recordings.
- public dataset for training and evaluating future models for insect detection and classification in camera recordings from natural environments.
- to evaluate different computing platforms, including edge devices, with respect to processing time.
- to connect the pipeline to insect ecology and conservation by demonstrating the proposed image processing pipeline in field-collected data.

## 2. Materials and methods

### 2.1. Processing pipeline

The *InsectDCT* tool first detects and localizes insects in the images, followed by taxonomic identification. The detection model is trained on a diverse set of images to improve robustness against false positives arising from small background objects that are visually similar to insects. A track-by-detection (TBD) approach was adopted due to its flexibility and because it does not require annotated tracking data. The pipeline can operate with or without tracking, depending on the selected time-lapse sampling interval. An overview of the proposed pipeline is illustrated in Figure 1 and elaborated below. The design prioritizes flexibility over computational efficiency and is based mainly on papers published by Bjerge et al. (2023b,c, 2025).

**Figure 1:**
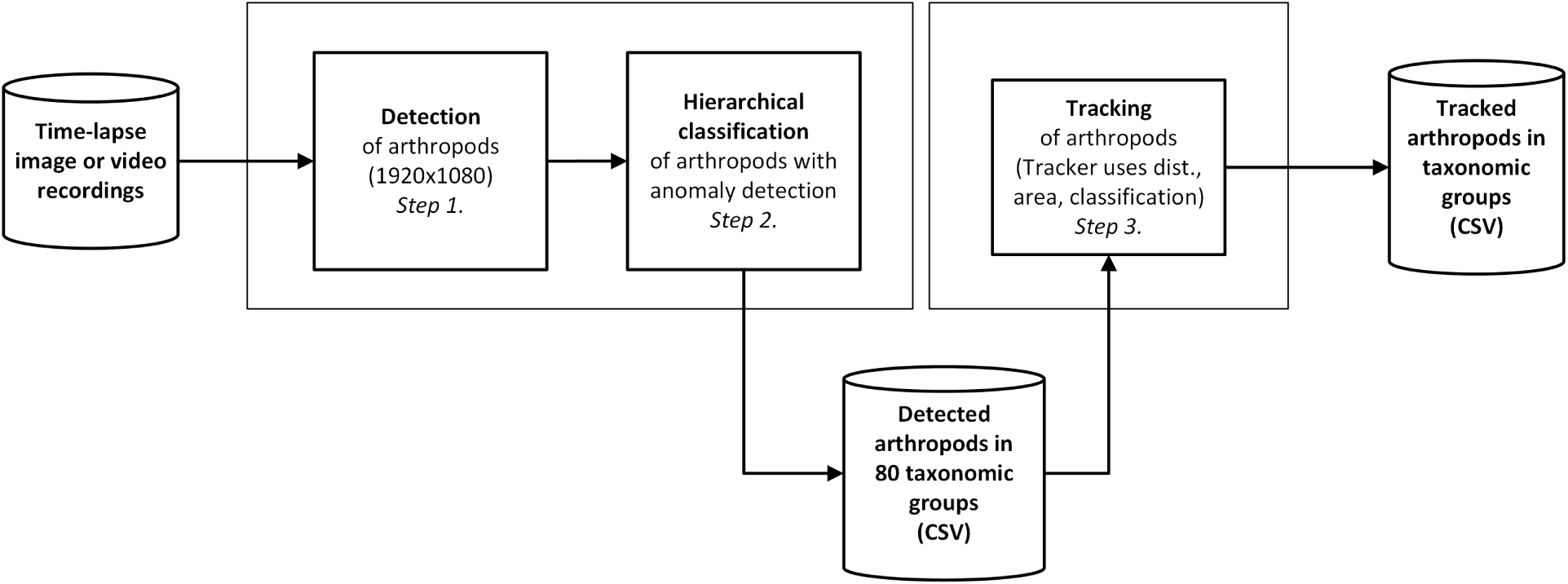
Processing pipeline to detect, classify and track insects from video or time-lapse recorded camera trap images. In step 1., the insects are detected and localized on resized images to 1920 × 1080 pixels using motion enhanced images and YOLO11. In step 2., insects are hierarchical classified into taxonomic ranks with 80 different taxonomic groups with anomaly detection of “unsure” insects. Localization and classification information of insect are used for the final tracking step 3. performed on time-lapse or video recorded with a sampling rate higher than 0.33 fps.

**Figure 2:**
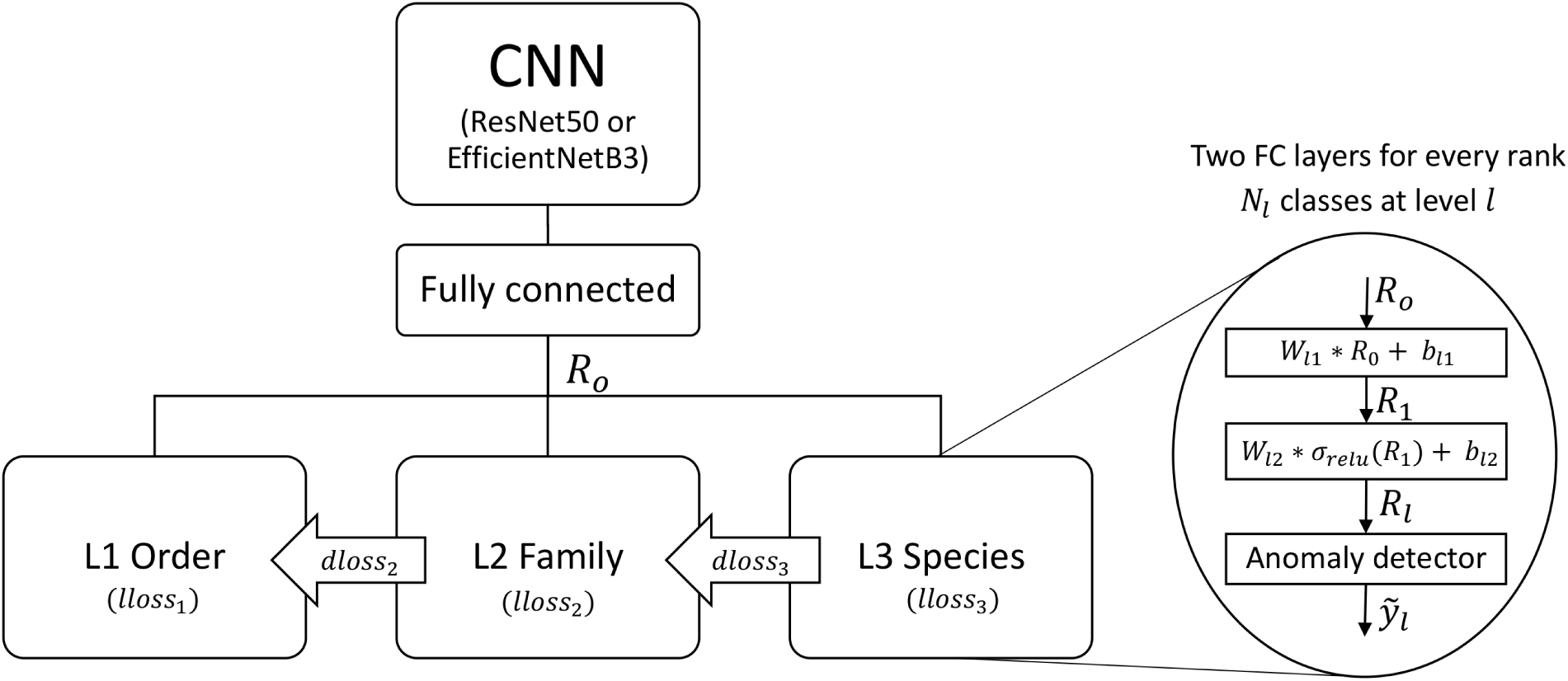
Network architecture with multitask learning to predict the three-level hierarchy. The multitask network shares the same backbone (ResNet50, EfficientNetV2 or ConvNext) with task branches at the end of parallel fully connected layers. A level (lloss) and dependency loss (dloss) function incorporates a level-dependent penalty to ensure hierarchy alignment. Figure from paper (Bjerge et al., 2023c).

**The first step** in our pipeline is to detect and localize insects in each image. This is achieved with an object detection model based on YOLO11 (Khanam and Hussain, 2024), trained on a dataset (see Section “Data collection for insect detection”) collected across diverse vegetation backgrounds and recorded with different cameras (see Tables 1 and 2). All images are resized to HD resolution (1920 × 1080 pixels), which reflects the typical resolution of the camera trap recordings used in the dataset. Medium and small versions of YOLO11 are trained on both standard color images and motion-enhanced images. Motion-enhanced images have previously been shown to improve recall (Bjerge et al., 2023b) by enabling the detection of more small insects (area 50 × 50 pixels), including those that are partially visible or visually cluttered with the background.

**Table 1:** A dataset of image samples was compiled from multiple sources to train insect detection and localization models across diverse vegetation backgrounds. The number of images, annotated arthropods, and empty background images without animals used for training and validation from each project is listed.

| Project name<br>(paper) | Images<br>train/val. | Arthropods<br>train/val. | Empty<br>train/val. | Plants | Cameras |
| --- | --- | --- | --- | --- | --- |
| **GreenRoof** (Bjerge et al., 2023a) | 9,385/2,453 | 9,441/3,151 | 980/37 | Sedum | Logitech |
| **GreenHouse** (Bjerge et al., 2023b) | 3,783/946 | 2,499/568 | 1,953/508 | Clover, Sea rocket | Logitech |
| **Arthropods** (Bjerge et al., 2024) | 6,901/525 | 4,770/766 | 2,654/299 | Sedum | Wingscapes |
| **PollWatch** | 2,867/720 | 468/118 | 2,488/625 | Mixed plants | Logitech |
| **MAMBO** | 4,313/1,163 | 5,725/1,635 | 806/206 | Sedum, Herbs | Wingscapes |
| **Orchard** | 3,157/798 | 2,705/696 | 703/173 | Sedum, Artificial | Pi V3 |
| **ACSHQ** (Serra-Marín et al., 2025) | 4,123/1,036 | 3,360/745 | 1,001/200 | >21 plant families | Pi HQ |
| Total | 34,529/7,641 | 28,995/7,763 | 10,612/2,048 | >30 plant families | 4 cameras |

**Table 2:**
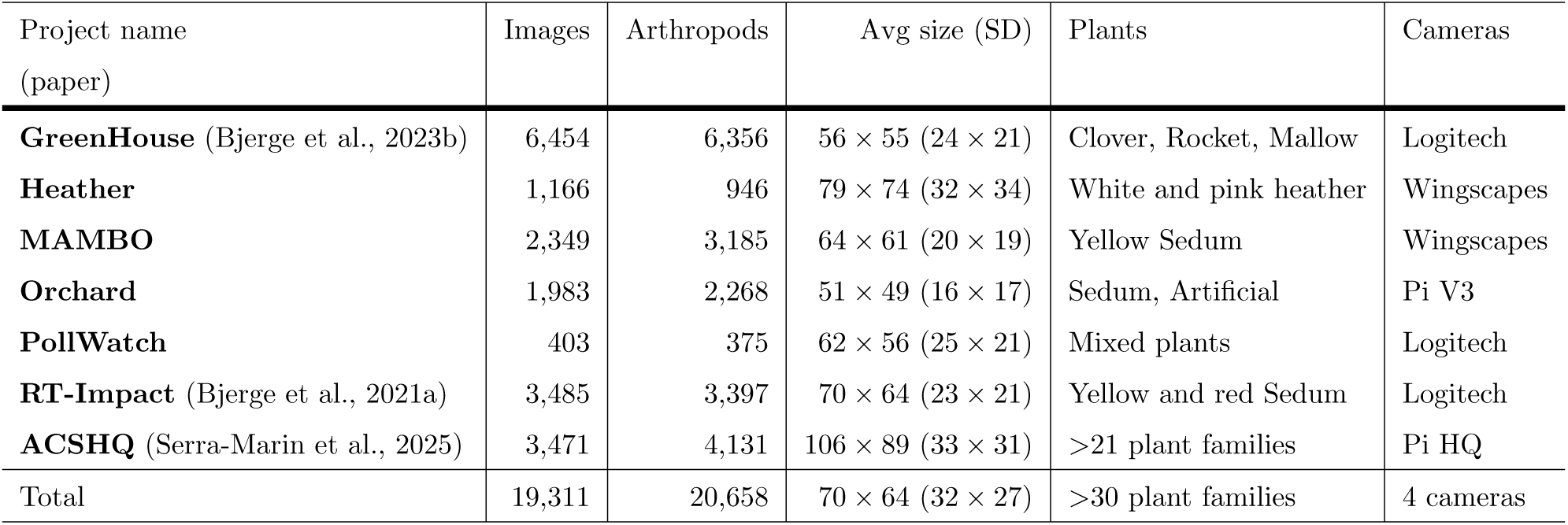
Test dataset of image samples was compiled from multiple sources to test insect detection and localization models across diverse vegetation backgrounds. The number of images, annotated arthropods, average pixel size and standard deviation (SD) used for testing from each project is listed.

**The second step** classifies the detected insects using a hierarchical classification model. The hierarchical classifier is trained on a dataset constructed from camera trap images collected in multiple locations and environments in Europe. The classifier architecture builds on the work of Bjerge et al. (2023c) and employs a multi-task learning framework combined with a dependency loss function that incorporates the taxonomic hierarchy. The hierarchical architecture consists of three taxonomic levels arranged in order, family, and genus/species. However, some of the levels also contain higher taxonomic ranks of animals seen in the footage. For each detection, the classifier outputs a label and an associated confidence estimate at each of the three taxonomic levels. An anomaly detection mechanism is integrated into the classifier to identify insects with class scores that are outside the distribution of the training dataset across taxonomic groups. These outliers may correspond to false-positive background detections, partially visible or blurry insects, or representatives of previously unseen insect classes or other animals. The output of the insect detector and hierarchical classifier is a list of insect detections with additional information about the camera trap, image, bounding box coordinates, confidence, class labels (Levels 1-3), date, and time.

**The third step** is based on track-by-detection using the bounding box area, the distance between detections, and the class label to create insect tracks. The tracker is based on the work published in Bjerge et al. (2025). A final list of insect tracks with information about the predicted insect taxon, species, confidence, size, date, arrival time, and duration seen by the camera is generated.

The source code for the pipeline is available on Github: https://github.com/kimbjerge/InsectDCT.

Each step in the pipeline is described below, with a summary of the algorithms described in original papers (Bjerge et al., 2023b,c, 2025).

#### 2.1.1. Insect detection and localization

Deep learning-based object detection methods rely primarily on spatial image information to extract features and identify object regions within an image. You Only Look Once (YOLO) (Redmon et al., 2016) is a one-stage object detector and is among the fastest available methods, making it well suited for processing millions of images or deployment on edge computing devices. In this work, YOLO11 (Glenn Jocher, 2020), with CSPDarknet53 as the backbone, was used.

Monitoring insects in their natural environments can be achieved using time-lapse cameras, where images or video are recorded at fixed intervals ranging from 60 to 0.033 seconds. Small moving objects are often easier to detect when temporal information is incorporated into the training of DL models. To address this, motion-informed detection, as proposed by Bjerge et al. (2023b), is used in this study to improve the detection of insect objects by exploiting temporal image sequences.

The motion-informed detection approach estimates movement by computing differences between consecutive frames in a time-lapse sequence. These differences are used to generate motion-enhanced images, which are subsequently used for both training and inference of the DL object detector. By modifying the standard RGB image format with motion information, existing object detectors can be used without architectural changes. This approach therefore enables the reuse of widely adopted convolutional neural network (CNN)–based object detectors such as YOLO (Redmon et al., 2016).

Three consecutive images from the time-lapse recording were used to generate each motion-enhanced image. The color images were first converted to grayscale and blurred using a Gaussian kernel of 5 × 5 pixels (image size: 1920×1080 pixels). These grayscale, blurred images were then used to compute a motion-vector 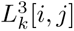 to modify the color RGB images. This process is summarized in equations eq. (1) and eq. (2).

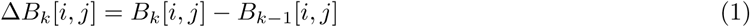

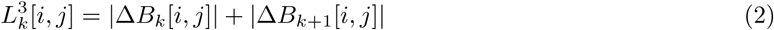

where [*i, j*] *∈* [1*..N* ] × [1*..M* ] are the pixel coordinates and *k ∈* N is the time index.

Here, the red channel is replaced with the motion-vector *L*^3^, whereas the blue channel is generated by combining the original red and blue channels with equal weighting (50% each). The result is blue/green images with red areas for where motion is detected. This approach makes it easier to find moving insects both visually and for the YOLO model. A detailed description of this process is provided in Bjerge et al. (2023b).

In this work, we trained four YOLO11 models (medium and small variants) using a comprehensive dataset consisting of both motion-enhanced images (MIE) and standard RGB images. This approach was adopted to develop a robust and generalizable model capable of detecting insects in various vegetation backgrounds and environmental conditions.

The performance of the insect detector and the localization model was evaluated in 5% of the training dataset, as well as in a separate test dataset containing images not used during training, including scenes with vegetation backgrounds previously unseen. The datasets for training, validation and testing are described in Section “Dataset for taxonomic classification”.

#### 2.1.2. Hierarchical taxon classifier with anomaly detection

The network architecture proposed by Bjerge et al. (2023c) is used and summarized in the following description. The architecture uses multitask learning by formulating taxonomy ranks as independent tasks to learn, which is implemented by modifying the output layer of a standard CNN. Specifically, the last layers are substituted with parallel fully connected (FC) layers for each taxonomy rank. Thus, a parameter sharing between all ranks while keeping rank-specific output layers. It is assumed that when more ranks are learned simultaneously, the more likely it is that the model will find a representation that captures all of the taxonomic ranks and dependencies.

The first part of the architecture is a CNN where we used ResNet (He et al., 2016), EfficientNetV2 (Tan and Le, 2021) or ConvNext (Todi et al., 2023). An input image *X* transformed by the network *N_CNN_* (*θ_CNN_* ), where *θ_CNN_*means the trainable parameters of the CNN network is given. The network output is viewed as a root representation *R*_0_,

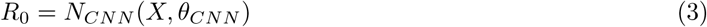

For each level *l* two FC layers and an activation function are added to perform a non-linear transformation of the input *R*_0_ to the level output *R_l_*.

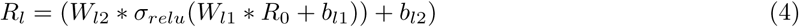

Here, *W_l_*_1_ and *W_l_*_2_ represent the weights and *b_l_*_1_*, b_l_*_2_ represent the biases for the independent representation of the FC layer. Dropout regularization is applied in the forward pass during training so that random neurons *W_l_*_1_ are disabled in the FC branches. Each rank is handled by the parallel FC layers to learn independent multiple levels of the taxonomy. This independent task representation is hierarchy-free, and the level dependency is handled by a combined loss function. Two types of losses are used, by combining a balanced softmax loss for each level and a hierarchical dependency loss function (Bjerge et al., 2023c). The balanced softmax (Jodelet et al., 2022) loss function is applied to compensate for the long-tailed distribution of the imbalanced dataset.

The images were cropped and resized to 128 × 128 or 224 × 224 pixels, which matches the dimensions used for a traditional CNN classifier (He et al., 2016). Training was performed using data augmentation, including image scaling, horizontal and vertical flip, and adding color jitter for brightness, contrast, and saturation. We selected a batch size of 64-256 for training our models, since it is faster to update and results in less noise than smaller batch sizes. The Adam optimizer with a fixed learning rate of 1.0 *·* 10*^−^*^4^ was chosen based on previously published experiments (Bjerge et al., 2023c). We have trained three CNN models to classify insects according to the hierarchical taxonomic groups defined by 80 classes described in Tables 3 and 4. CNN models were fine-tuned using pre-trained weights from ImageNet (Russakovsky et al., 2015).

**Table 3:**
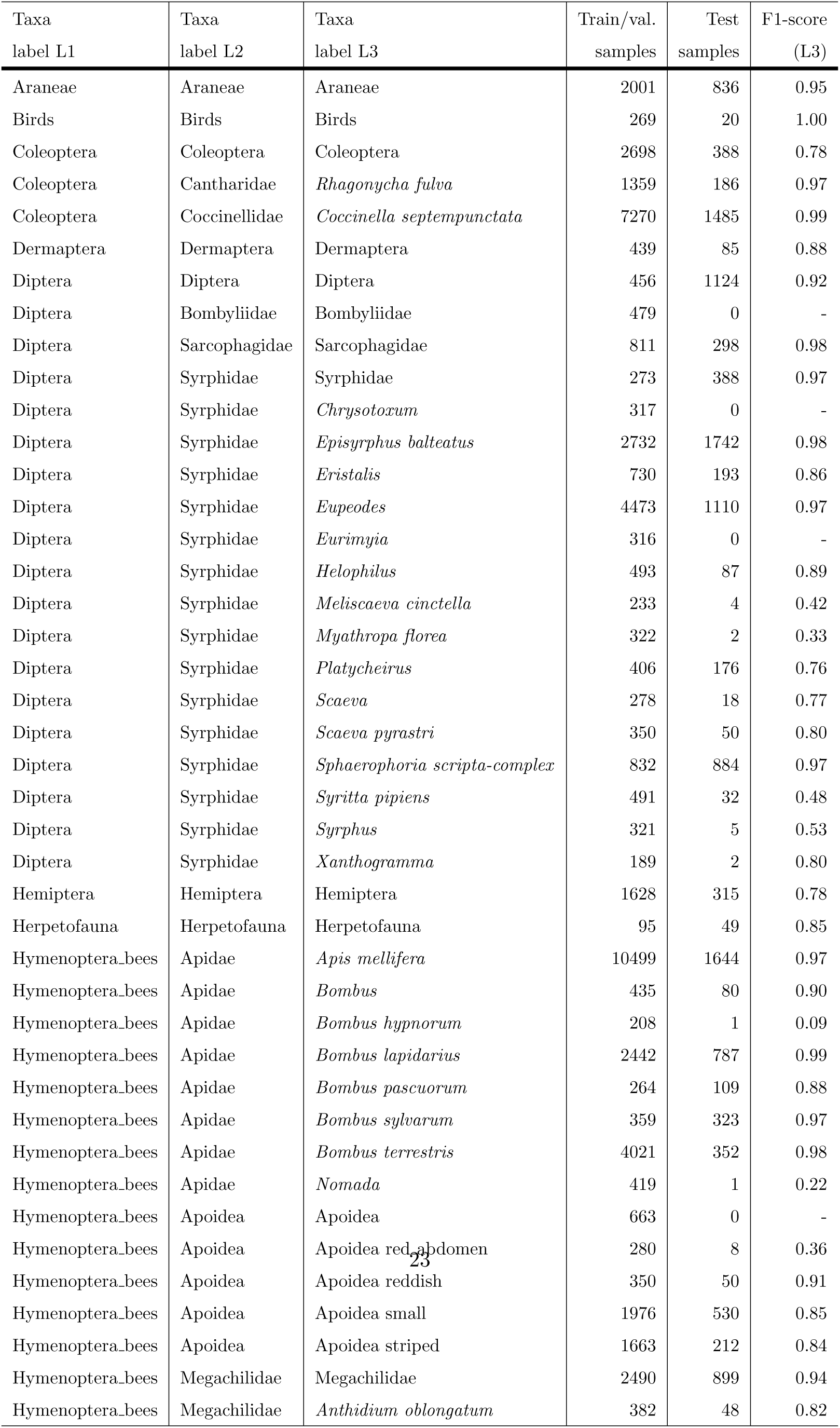
Dataset (Part 1): Image samples used for training and testing a taxonomic classifier. Class names and the number of samples are listed for labels at hierarchical levels 1, 2, and 3 (L1, L2, and L3). The F1-score of the best-performing ConvNeXt-Base model is reported for each taxon at L3, evaluated on the test samples.

**Table 4:**
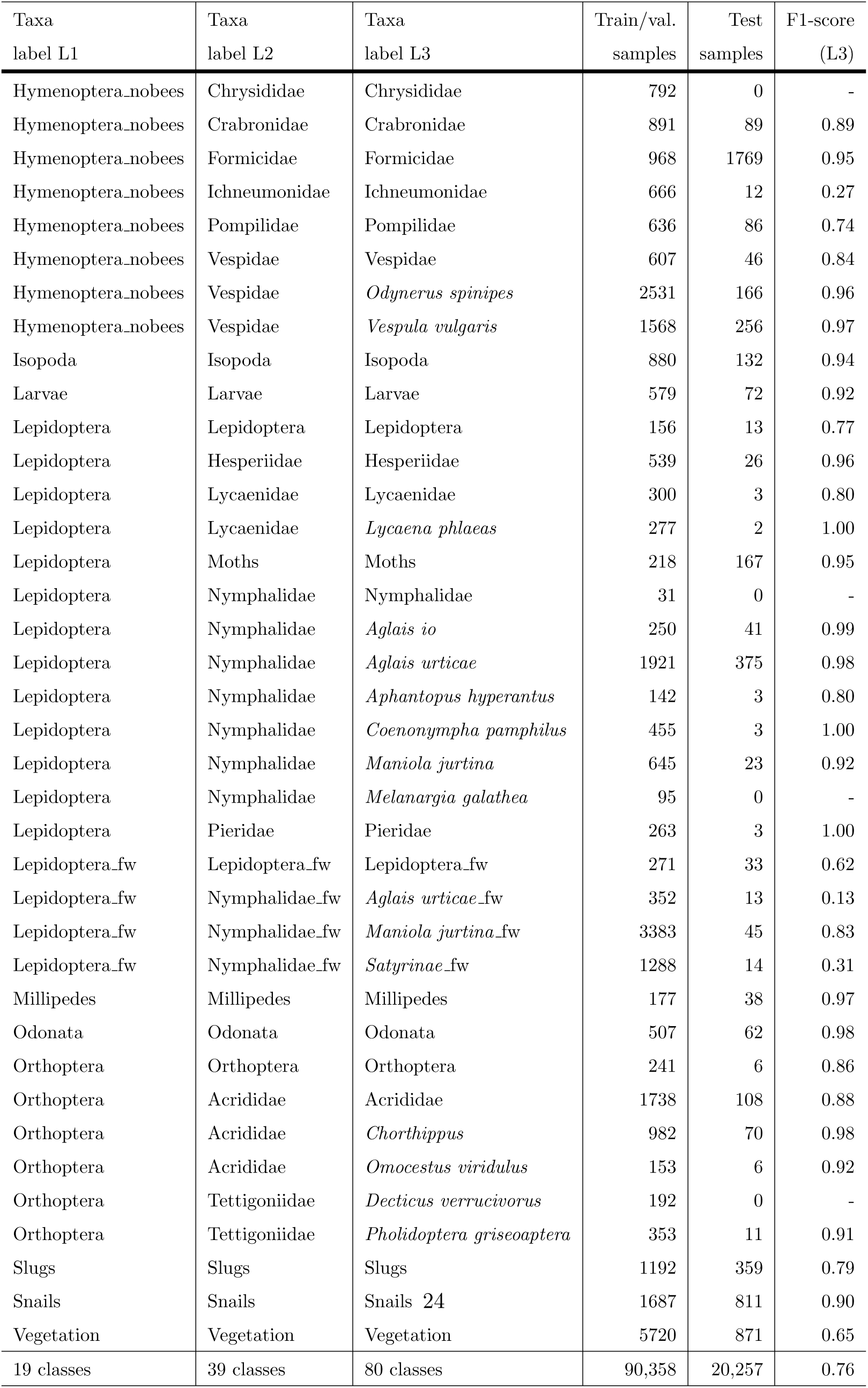
Dataset (Part 2): Image samples used for training and testing a taxonomic classifier. Class names and the number of samples are listed for labels at hierarchical levels 1, 2, and 3 (L1, L2, and L3). The F1-score of the best-performing ConvNeXt-Base model is reported for each taxon at L3, evaluated on the test samples.

##### Anomaly detection

The methodology of out-of-distribution detection (Bulusu et al., 2020) and threshold-based anomaly tagging is used to identify instances of “anomalies” such as uncertain classifications. In our application, these instances may manifest as debris, obscured or partially visible insects, or those exhibiting blurriness, characteristics that are not represented in the insect taxon training dataset.

Often, softmax is the last layer in a classification neural network, where the maximum value determines the predicted class. Here, we instead analyze the output distribution without the softmax layer to determine the anomalies and predict the classes. The distribution of the output *x* for each predicted class *j^th^* follows a normal distribution 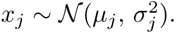.

An example of the output distribution is shown in Figure 3 which is generated on the sample training dataset (Coleoptera) for corrected classified input. If the output value *x_j_* is below a threshold of *th* = *µ−*3.0*σ*, we label the input as anomaly. Consequently, when new unknown inputs are presented for the trained network and the output lies below the threshold, it will be classified as an “uncertain” prediction. The threshold is set to ensure that fewer than 1% of correctly classified inputs are marked “uncertain”. However, thresholds between *µ −* 2.0*σ* and *µ −* 6.0*σ* can also be selected, depending on the desired strictness of the anomaly detector and the backbone classifier.

**Figure 3:**
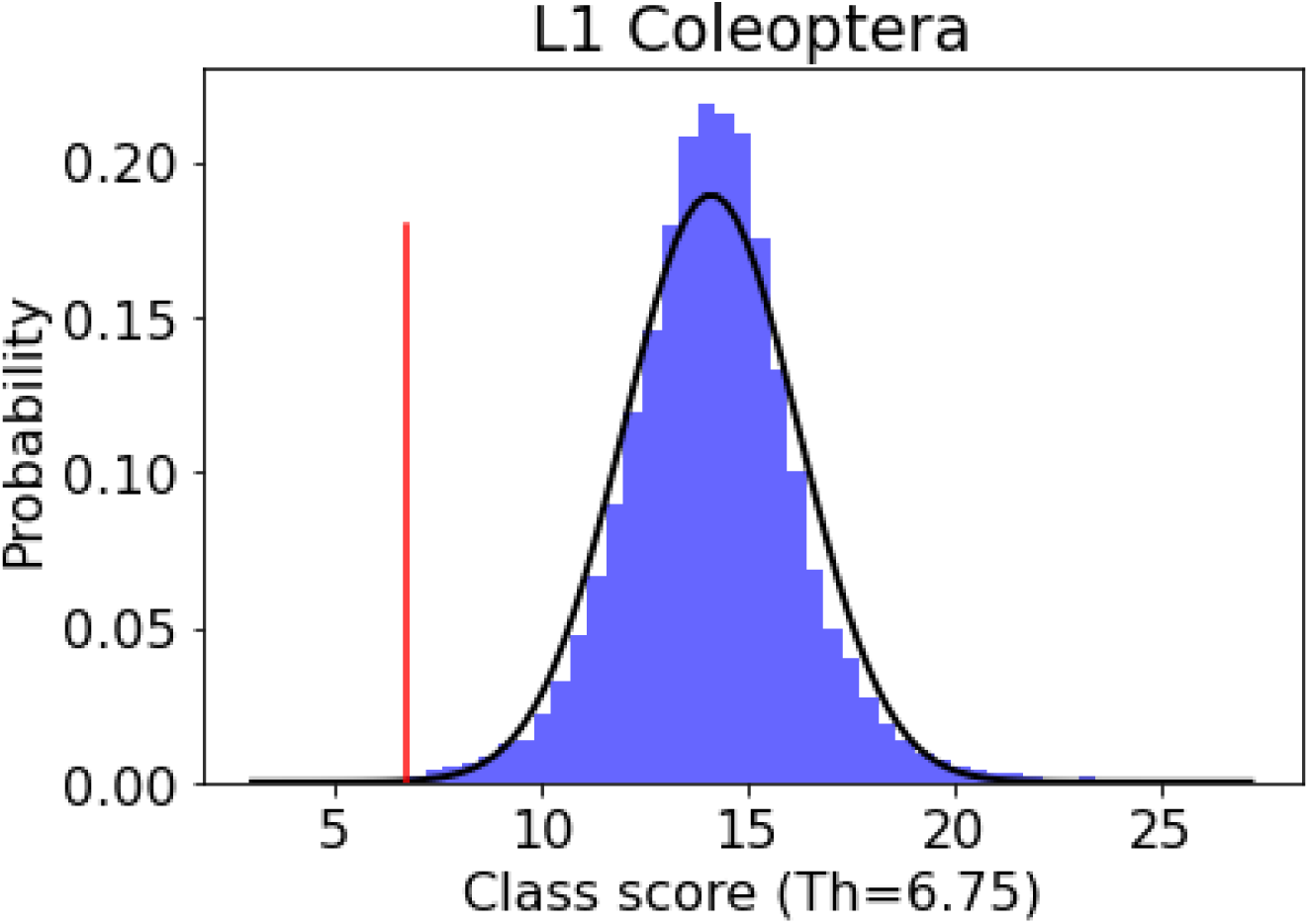
Probability density function for the output scores for Coleoptera and the chosen threshold for uncertain anomalies.

If a classification is correct at a higher taxonomic level, it is still considered valid and useful, even if it remains “unsure” at lower levels of the hierarchy. A classification is also labelled “unsure” if the predicted labels across the three levels are not taxonomically consistent.

This threshold-based filtering is applied to refine classifications to the highest reliable taxonomic resolution, with pollinators identified at the genus or species level whenever sufficient confidence is achieved. By incorporating anomaly detection, the classifier can assign specimens to higher taxonomic ranks when predictions at lower levels are uncertain. For example, an individual classified as “unsure” at levels 2 or 3 could still be reliably identified as belonging to the order Hymenoptera at level 1. This hierarchical approach preserves useful taxonomic information while reducing the risk of overconfident misclassification at finer taxonomic resolutions.

Finally, the output scores *x_j_* are assigned a probability *F* (*x_j_*) by estimating the cumulative distribution function as the integral of the probability density function given by. This approach provides a statistical measure of the predictions, in contrast to the softmax confidence score.

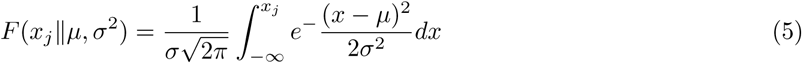

#### 2.1.3. Tracking

The tracking algorithm proposed by Bjerge et al. (2021a) is used and described briefly below. The Hungarian algorithm was selected to determine the optimal assignment within a given cost matrix, enabling the association of detections across successive images in an efficient and globally optimal manner. In this application, the cost matrix should represent the probability that an insect in the previous image had moved to a given position in the current image. The cost function was defined as the weighted cost of distance and area of the matching bounding boxes in the previous and current images. The Euclidean distance *D* between the central position (*x, y*) in the two images was calculated as follows.

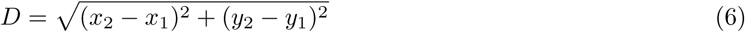

This distance was normalized according to the diagonal of the image *I*:

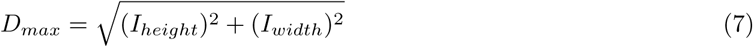

The cost of the area was defined as the cost between the area *A* of the bounding boxes:

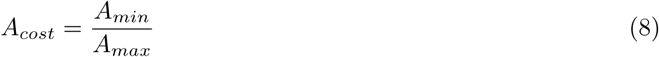

A final cost function in Equation (9) was defined with a weighted distance cost *W_d_* and a weighted area cost *W_a_*.

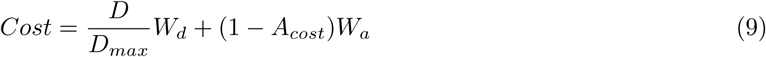

A cost threshold was established to determine whether successive insect detections should be associated. Subsequently, a track was established, stipulating a minimum of two detections per track. The weights *W_d_* and *W_a_* are set to 1.3 and 0.7, respectively, giving greater importance to the distance between detections than to differences in bounding-box area when assigning detections to tracks. These values were adopted from the original proposal of the tracker published in (Bjerge et al., 2021b). For each track, information such as the start date and time, duration, number of detections, and average size was recorded. Of particular importance was the recording of the predominant insect taxon, along with the accuracy of its classification. Ultimately, a track was considered valid when *>*50% of the detections corresponded to the predominant classification and comprised at least three detections. The tracker can be configured to incorporate classification results when forming tracks, ensuring that only classifications belonging to the same taxonomic group are associated. This optional extension is particularly beneficial when the classification results are sufficiently accurate.

### 2.2. Data collection and datasets for pipeline training and evaluation

We collected time-lapse and video recordings from several monitoring sites using different cameras, including Logitech C922 web cameras (2MP), Wingscapes Time-lapse Pro Bird cameras (10MP), Raspberry Pi V3 cameras (2MP), and Raspberry Pi High Quality (HQ) cameras (HD video). Published studies by Bjerge et al. (2021a, 2023a,b, 2024); Serra-Marin et al. (2025) include insect recordings from environments containing more than 24 different plant families and even more species, and a subset of these recordings was incorporated into our datasets. A balanced dataset was constructed by selecting 4,000-10,000 images from each source, representing various backgrounds and camera types, to train a generalized insect detector. In addition, four other projects (listed below) contributed recordings obtained with various cameras and plant types, including *Sedum*, mixed plants and flowers, herbs, and artificial flowers. For each project, between 10 and 30 cameras were used to collect images during the monitoring season. These additional projects, provide time-lapse recordings of a single flowering season (May-September).

- **Heather** - Wingscapes Pro camera recordings of heather plants (Denmark, 2019)
- **PollWatch** - Logitech C922 camera recordings of mixed plants and flowers (Denmark, 2020)
- **MAMBO** - Wingscapes Pro camera recordings of *Sedum* and herb plants (Denmark, France, Germany, Malta, Netherlands, United Kingdom, 2024)
- **Orchard** - Raspberry Pi V3 camera recordings of *Sedum* plants (Germany, 2025)

Two datasets were created: one for training a generalized insect detector model and another model for taxonomic classification of detected insects at the order, family, and species levels. The aim is to train one deep learning (DL) model to detect insects and a second DL model to classify the detected insects into taxonomic groups. The training and validation datasets may contain temporally adjacent images, as the split was performed at the image level. However, the test datasets used to evaluate both the insect detector and the hierarchical classifier consist exclusively of recordings from different dates than those used for training and validation. Consequently, the test images are not temporally adjacent to the images in the training or validation sets, and some images even have different backgrounds.

The classification model is trained on the recordings described in this paper, but can be extended to include additional taxonomic groups with additional data provided. The two datasets for detection and taxonomic classification are described below and are available at: https://doi.org/10.5281/zenodo.21154489

#### 2.2.1. Dataset for insect detection

With time-lapse images, a substantial proportion of these images may not contain any arthropods, resulting in false-positive detections of plant structures and other background elements (Bjerge et al., 2024). Therefore, an insect detection dataset was constructed through an iterative process of progressively incorporating images that contain diverse insect species and vegetation backgrounds with false-positive detections. An initial dataset was formed by combining datasets published by Bjerge et al. (2023a,b, 2024), and a DL model was trained to identify additional images containing arthropods from other projects with recorded data. Images containing insect detections were manually reviewed to remove false positives and identify missed detections by inspecting each image and correcting the corresponding bounding boxes using a labeling tool. This procedure was repeated for each project, with new DL models trained at every iteration as additional images were incorporated into the dataset. With each iteration, the number of false-positive detections was progressively reduced, improving the overall robustness of the detection model.

The final dataset, summarized in Table 1, comprises 42,170 images that contain arthropods captured against challenging backgrounds, including more than 30 different plant families. In total, the dataset includes 36,758 annotated arthropods and other animals identified in the images. Certain animals, such as slugs and snails, were excluded and therefore not annotated. The dataset consists of both color images and motion-enhanced images.

To evaluate the performance of the insect detector, a test dataset was created that includes two new projects (new locations and plants) and images from the same projects but from camera backgrounds not used in the training and validation dataset. The test dataset comprises recordings from days that were not included in the training or validation datasets, enabling the evaluation of the model’s ability to transfer to new environments without requiring the training of a project-specific detector.

The final test dataset, summarized in Table 2, comprises 19,311 images that contain insects captured against challenging backgrounds, including backgrounds with new plant families (different species of heather plants). Example images from the datasets are presented in the Supporting information.

#### 2.2.2. Dataset for taxonomic classification

For the classification of detected insects, two datasets were created to train and evaluate hierarchical classification models. The datasets were organized into three taxonomic levels: order at level 1, family at level 2, and genus/species at level 3. The datasets were structured on the basis of the taxonomic groups of insects observed in the camera recordings. Figure 4 presents example images of selected pollinators grouped at the species level. However, many pollinator taxa could only be reliably classified or annotated at the genus level. This also explains why Level 3 combines genus- and species-level classifications rather than separating them into two distinct levels.

**Figure 4:**
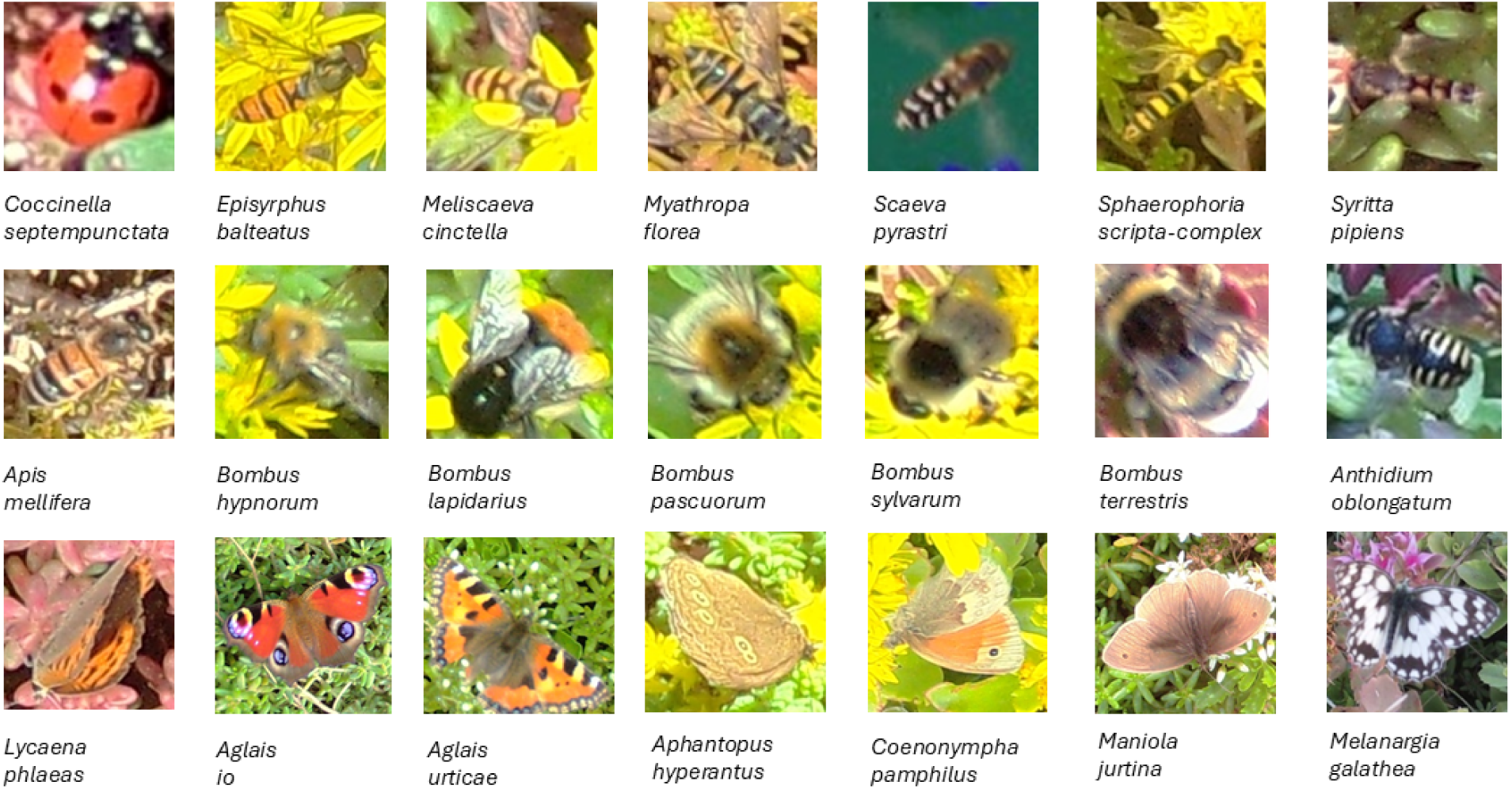
Shows example images of selected pollinators grouped at the species level, covering the families Coccinellidae, Syrphidae, Apidae, Megachilidae, Lycaenidae, and Nymphalidae.

Most of the recordings, were collected from plots, dominated by Sedum plants, using images from projects that were also included in the detection datasets. The training dataset was compiled from recordings obtained in the projects: **GreenRoof** (Bjerge et al., 2023a), **Arthropods** (Bjerge et al., 2024), **MAMBO**, and **Orchard**. These projects were selected because they contained a wide diversity of insect species, many of which had already been annotated in previous studies. The training data were further supplemented with GBIF images corresponding to the species identified in the camera recordings. This dataset was divided into 95% for training and 5% for validation.

The test dataset was created using recordings from some of the same projects but from different camera locations, including **RT-Impact** (Bjerge et al., 2021a), **Arthropods** (Bjerge et al., 2024), **MAMBO** and **Orchard**. This dataset consists of recordings from dates that were not represented in the training or validation data, thereby ensuring that the reported classifier performance is evaluated on temporally separated observations. The purpose of the test dataset was to evaluate the models within the same insect domain while assessing generalization across different camera trap locations. The test dataset contains many of the same species as the training dataset; however, some species are not represented, representing a challenge to model generalization.

Tables 3 and 4 present the complete training and test datasets, comprising 19 taxa at level 1, 39 taxa at level 2, and 80 taxa at level 3. Some classes represent broader taxonomic categories, including animals and vegetation at higher taxonomic ranks. The classes were defined based on the organisms detected in the images, including false-positive detections of vegetation.

Special Apoidea groups were introduced at level 2 for bees that could not be reliably identified at the genus or species level due to limited image quality or insufficient visual detail. In several cases, taxa at level 3 were further divided into morphogroups such as Apoidea red abdomen, small, and striped, where image size and quality were insufficient to reliably identify species-level characteristics. Only pollinator groups Syrphidae, Hymenoptera bees, and Lepidoptera were consistently annotated at the genus or species levels, reflecting their primary relevance to the purpose of the classifier. Other groups were identified only at higher taxonomic levels and, therefore, assigned the same label across lower levels of the hierarchy. For example, specimens classified as the order Hemiptera at level 1 are also labeled as Hemiptera at levels 2 and 3.

The average size of the insect images is 227 × 224 with a standard deviation of 235 × 234 pixels. However, as shown in Figure 5, the range of image sizes was wide, with GBIF images generally much larger than those captured by camera traps. Consequently, images recorded using insect camera traps contain less visual information, which can affect classification performance.

**Figure 5:**
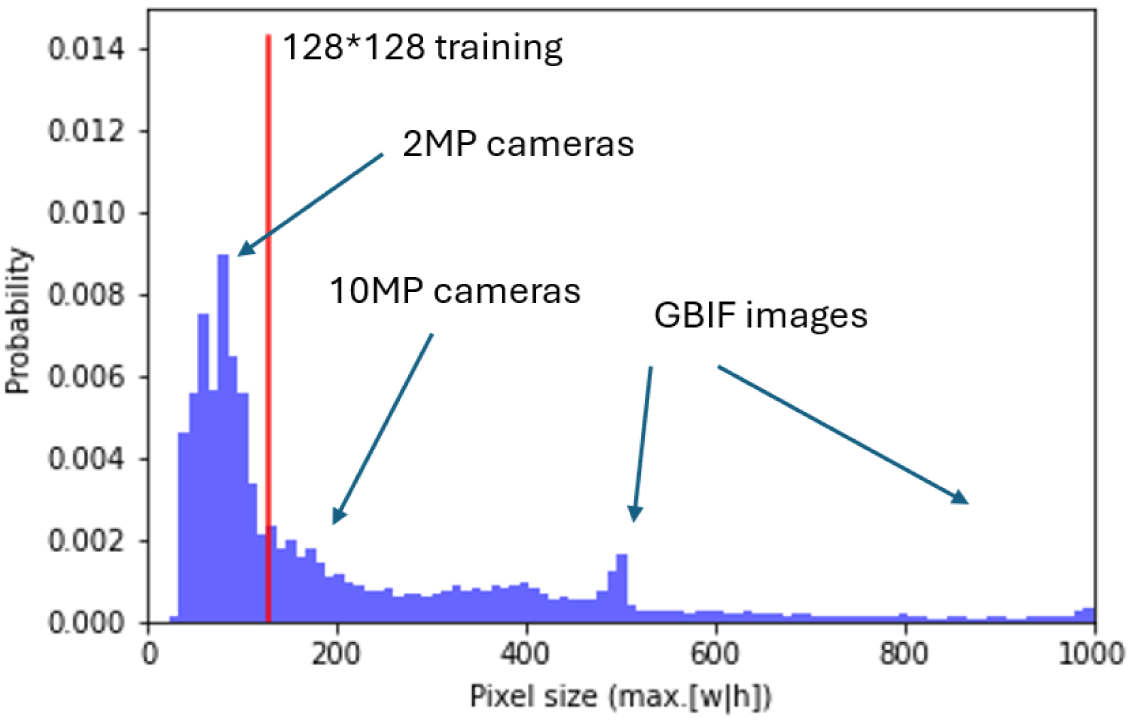
The distribution of insect image sizes in the training dataset. During training, all images are resized to 128 × 128 pixels. Insects captured in the 10 MP recordings and GBIF images are generally larger than those from the 2 MP recordings.

#### 2.2.3. Dataset for tracker performance evaluation

To evaluate tracker performance under challenging conditions, recordings with low temporal sampling rates were selected for the tracking evaluation. In total, eight recording days from different camera locations were used. Six days were recorded at different **Orchard** camera locations using two-second time-lapse intervals (0.5 fps), while two days were recorded with the **RT-Impact** camera using three-second intervals (0.33 fps). For each recording day, a period of high insect activity was selected, resulting in evaluation sequences containing between 100 and 440 frames. Insects detected during these periods were manually curated to establish ground-truth (GT) tracks, which were then used to evaluate the multiple-object tracking (MOT) performance of the proposed tracker.

The evaluation included several tracking metrics. Identity switches (IDSW) were recorded when a predicted track ID was incorrectly assigned to a different GT track. Track fragmentation events (FM) were measured by counting gaps of less than 10 s within a GT track that resulted in the insect being represented by multiple predicted tracks. In addition, Multiple Object Tracking Accuracy (MOTA) Bernardin and Stiefelhagen (2008); Milan et al. (2016) and Identification F1 Score (IDF1) Ristani et al. (2016) were calculated to provide complementary measures of overall tracking accuracy and identity preservation.

## 3. Results

### 3.1. Insect detection and localization

Precision, recall, F1-score, and mAP50 for the detection and localization stage of the pipeline are presented in Table 5 evaluated in the validation dataset. The larger YOLO11m model achieves the highest F1-score; however, the improvement is marginal considering the increased model size, with YOLO11m having 20.1M parameters compared to 9.4M for YOLO11s. Models trained on motion-enhanced images show improved F1-scores, with particularly higher values of mAP50 and mAP50-95. In general, the validation results in Table 5 indicate that there are no substantial differences between the models, as all perform well.

**Table 5:** Validation results for insect detection and localization on 5% of the training dataset with 7,641 images and 7,763 labelled insects.

| Model | Image | Parameters (M) | Precision | Recall | F1-score | mAP50 | mAP50-95 |
| --- | --- | --- | --- | --- | --- | --- | --- |
| YOLO11s | RGB | 9.4 | 0.914 | 0.924 | 0.919 | 0.945 | 0.658 |
| YOLO11m | RGB | 20.1 | 0.909 | 0.933 | 0.921 | 0.946 | 0.654 |
| YOLO11s | MIE | 9.4 | 0.923 | 0.934 | 0.928 | 0.956 | 0.682 |
| YOLO11m | MIE | 20.1 | 0.923 | 0.935 | 0.929 | 0.951 | 0.711 |

Table 6 presents the precision, recall, F1-score and mAP50 of the insect detector evaluated across seven different sources included in the test dataset, each comprising collected and annotated images with various backgrounds of plants and insects. The YOLO model trained with motion-enhanced images demonstrates a notable improvement in performance, achieving an average F1-score of 0.883 compared to 0.806 for the model trained on standard RGB images. In addition, the standard deviation across datasets is lower for the model trained on motion-enhanced images, indicating a more consistent performance. The improvement is particularly evident in the recall, which increases from an average of 0.753 to 0.853. This indicates that the model trained on motion-enhanced images detects approximately 10% more insects compared to the standard model trained on RGB images. Compared to the validation results in Table 5, the performance decreases in the test dataset, as it contains images that were not used during training. However, the reduction in F1-score is only 4.6% for the model trained with motion-enhanced images, compared with 11.5% for the model trained on standard RGB images.

**Table 6:** Test results for insect detection and localization using YOLO11m on the test dataset comprising 19,311 images and 20,658 labeled insects. The YOLO11m model is trained with color (RGB) images or motion-enhanced images (MIE). The table reports the average performance across the seven project datasets, along with the corresponding standard deviation (SD).

| Project | Image | Precision | Recall | F1-score | mAP50 | mAP50-95 |
| --- | --- | --- | --- | --- | --- | --- |
| **GreenHouse** | RGB | 0.905 | 0.549 | 0.683 | 0.719 | 0.455 |
| **GreenHouse** | MIE | 0.902 | 0.747 | 0.817 | 0.886 | 0.565 |
| **Heather** | RGB | 0.930 | 0.801 | 0.860 | 0.888 | 0.674 |
| **Heather** | MIE | 0.896 | 0.901 | 0.898 | 0.953 | 0.788 |
| **MAMBO** | RGB | 0.812 | 0.699 | 0.751 | 0.810 | 0.670 |
| **MAMBO** | MIE | 0.908 | 0.879 | 0.893 | 0.955 | 0.846 |
| **Orchard** | RGB | 0.844 | 0.859 | 0.851 | 0.903 | 0.734 |
| **Orchard** | MIE | 0.890 | 0.891 | 0.890 | 0.949 | 0.791 |
| **PollWatch** | RGB | 0.853 | 0.665 | 0.748 | 0.794 | 0.621 |
| **PollWatch** | MIE | 0.899 | 0.795 | 0.844 | 0.902 | 0.723 |
| **RT-Impact** | RGB | 0.939 | 0.807 | 0.868 | 0.899 | 0.712 |
| **RT-Impact** | MIE | 0.961 | 0.926 | 0.943 | 0.979 | 0.822 |
| **ACSHQ** | RGB | 0.880 | 0.891 | 0.883 | 0.926 | 0.656 |
| **ACSHQ** | MIE | 0.924 | 0.868 | 0.895 | 0.947 | 0.790 |
| Average (SD) | RGB | 0.880 (0.043) | 0.753 (0.112) | 0.806 (0.072) | 0.848 (0.070) | 0.656 (0.090) |
| Average (SD) | MIE | 0.911 (0.023) | 0.858 (0.059) | 0.883 (0.038) | 0.939 (0.030) | 0.761 (0.087) |

### 3.2. Hierarchical taxon classifier with anomaly detection

A total of 5% of the training dataset was reserved for validation and early stopping was applied after 45, 72, and 66 epochs, respectively. This relatively small percentage was chosen because the validation dataset was used solely for early stopping and not for the final model performance evaluation. The three models were then evaluated on the test dataset, with the results summarized in Table 7.

**Table 7:** Performance results for three hierarchical classifiers using ResNet50V2, EfficientNetV2-S, and ConvNeXt-base (CNB) backbones. Precision, recall, and F1-score are reported for each level (L1, L2, and L3) of the taxonomic hierarchy, evaluated on the test dataset. The backbone achieving the highest F1-score at each level is highlighted in bold. The number of model parameters and classes at each level is listed.

| Model-Level | Parameters<br>(M) | Classes | Precision | Recall | F1-score<br>(macro) | F1-score<br>(micro) |
| --- | --- | --- | --- | --- | --- | --- |
| ResNet-L1 | 25.6 | 19 | 0.799 | 0.904 | 0.831 | 0.912 |
| EfficientNet-L1 | 21.6 | 19 | 0.833 | 0.921 | 0.860 | 0.925 |
| CNB-L1 | 88.5 | 19 | 0.897 | 0.926 | **0.903** | **0.948** |
| ResNet-L2 | 25.6 | 39 | 0.768 | 0.880 | 0.783 | 0.907 |
| EfficientNet-L2 | 21.6 | 39 | 0.790 | 0.894 | 0.804 | 0.921 |
| CNB-L2 | 88.5 | 39 | 0.840 | 0.916 | **0.843** | **0.939** |
| ResNet-L3 | 25.6 | 80 | 0.698 | 0.843 | 0.693 | 0.884 |
| EfficientNet-L3 | 21.6 | 80 | 0.727 | 0.850 | 0.728 | 0.900 |
| CNB-L3 | 88.5 | 80 | 0.758 | 0.889 | **0.759** | **0.922** |

The F1-score increases at higher levels of the taxonomic hierarchy. The macro F1-score is higher than the micro F1-score across all models, reflecting the long-tailed and imbalanced distribution of both the training and test datasets. ConvNeXt-base achieves the best overall performance, followed by EfficientNetV2-S and ResNet50V2. However, EfficientNetV2-S has the lowest number of parameters, which makes it more suitable for applications where computational efficiency and inference speed are important.

The best-performing ConvNeXt-base model achieves average F1-scores of 0.90 at level 1 (L1), 0.84 at level 2 (L2), and 0.76 at level 3 (L3). Tables 3 and 4 show that the relatively low average F1-score of 0.76 is mainly driven by seven classes with very few test samples (1–13 samples per class). In contrast, the micro F1-score is substantially higher (0.92), indicating that the limited sample size in these classes introduces increased uncertainty in the overall evaluation.

The seven taxa with few test samples and correspondingly low F1-scores at level L3 were *Meliscaeva cinctella* (0.42), *Myathropa florea* (0.33), *Bombus hypnorum* (0.09), *Nomada* (0.22), Apoidea red abdomen (0.36), Ichneumonidae (0.27) and *Aglais urticae* fw (0.13). Taxa without dedicated test samples were evaluated using the 5% validation dataset and achieved high F1-scores, including Bombyliidae (0.98), *Chrysotoxum* (0.97), *Eurimyia* (0.94), Apoidea (0.74), Chrysididae (0.99), Nymphalidae (1.0), *Melanargia galathea* (1.0) and *Decticus verrucivorus* (0.9). However, these validation scores are likely optimistic, as the performance evaluated on the independent test datasets was generally lower.

The metrics reported in L2 and L3 allow for predictions within the same taxonomic hierarchy. For example, a prediction of *Bombus* is considered correct if the true label is *Bombus lapidarius*. This hierarchical correction is not applied in Figure 6, which illustrates how varying the threshold of anomaly detection affects performance. As more predictions are labeled as “unsure”, the F1-score increases. A selected anomaly threshold value of 5.5 results in approximately 4%–8% of the predictions being labeled as “unsure”. When a prediction is labeled as “unsure” in L2 or L3, the corresponding prediction is used instead at the next higher taxonomic level.

**Figure 6:**
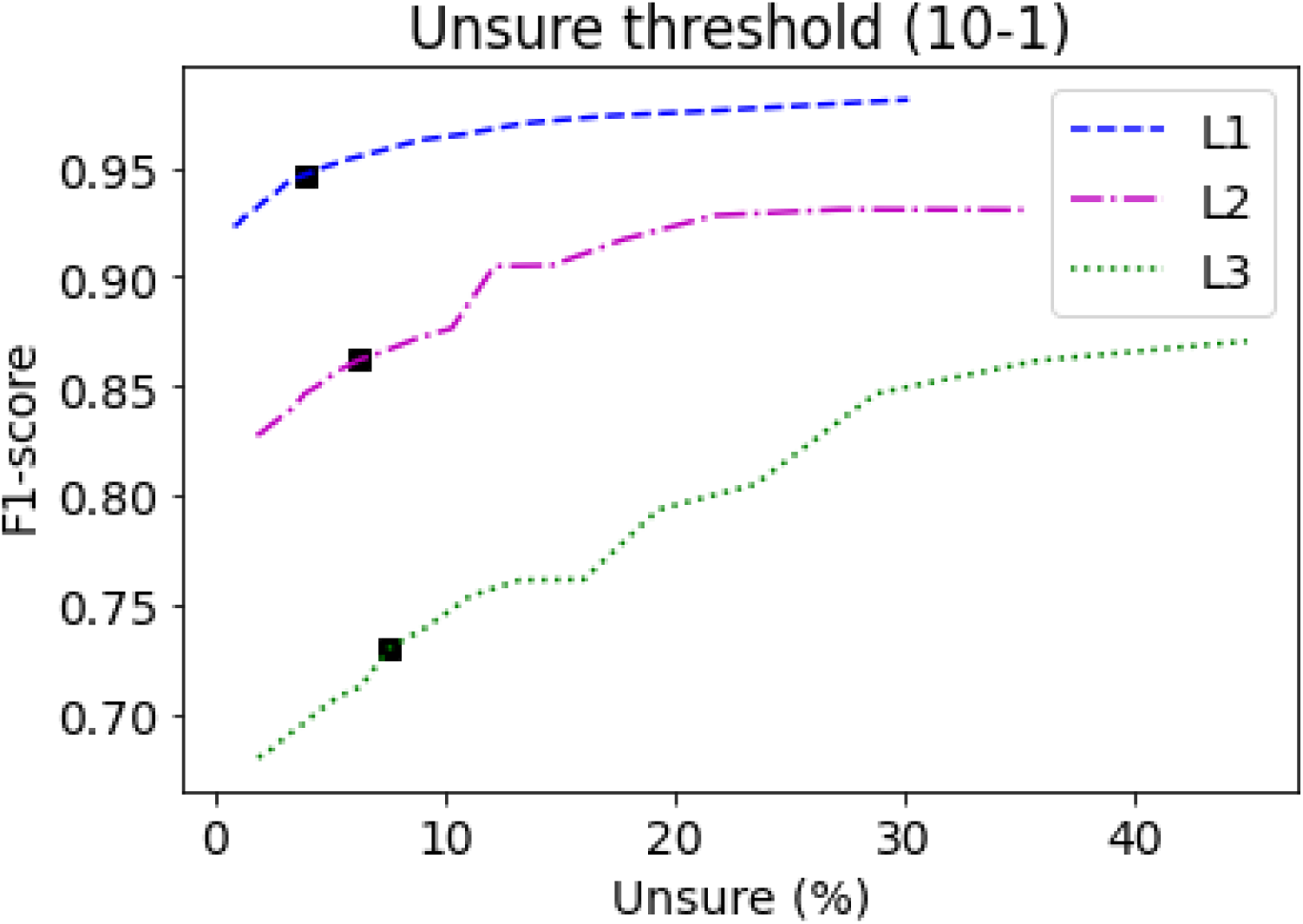
The F1-score at each taxonomic level for different unsure thresholds. As the uncertainty threshold increases, the proportion of classifications labelled as uncertain also increases, while the F1-score improves. The selected threshold value of 5.5 is indicated by a bold square using ConvNeXt-Base as the backbone.

The confusion matrix for level 1, shown in Figure 7, indicates that the classes Lepidoptera and Lepidoptera fw (folded wings) are frequently misclassified. In addition, 21% of slugs are predicted to be snails. Approximately 32% of vegetation samples are classified as “unsure”; however, this is acceptable, as the primary objective is to eliminate false-positive background detections including vegetation.

**Figure 7:**
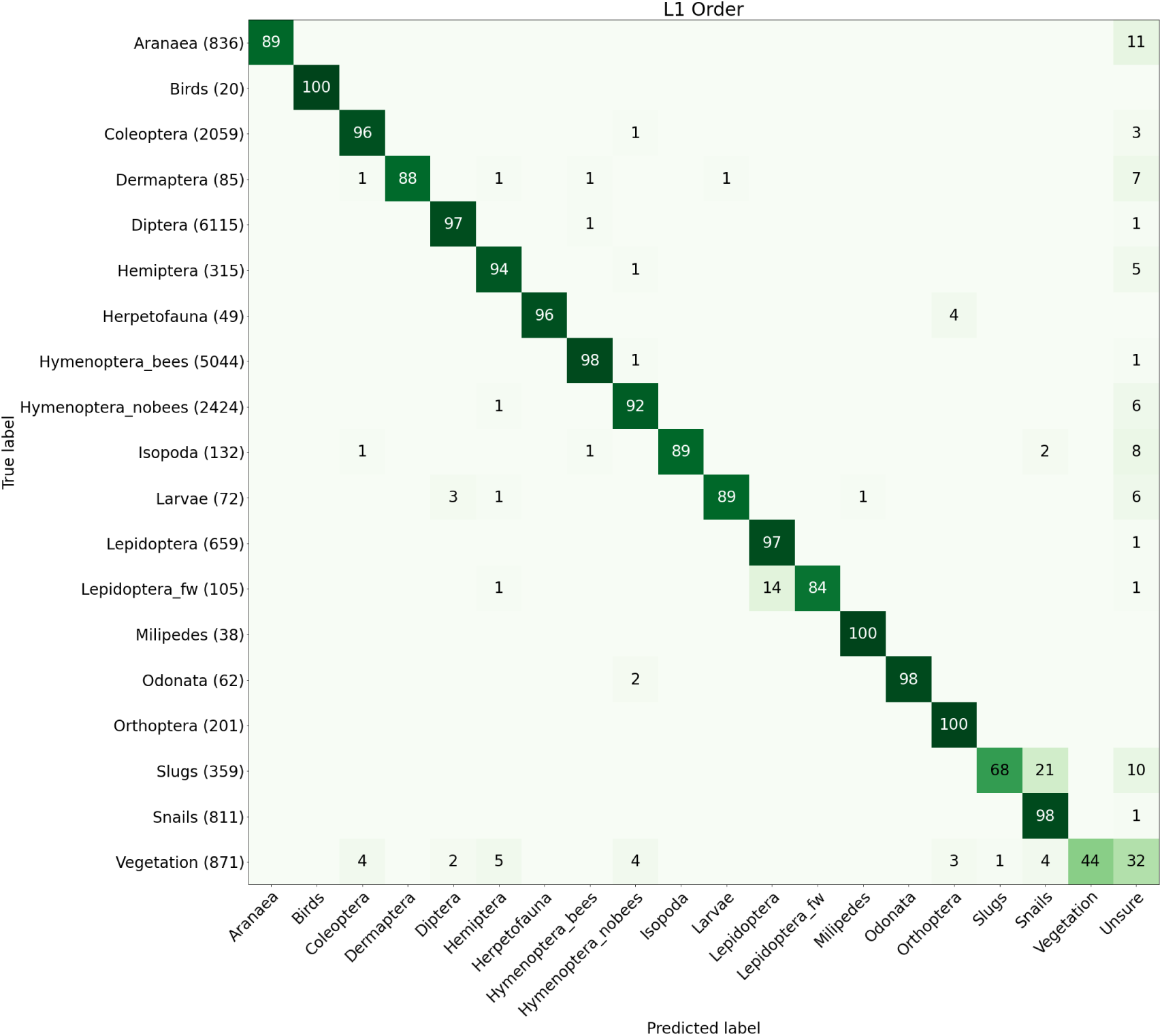
Confusion matrix for level 1 predictions on the test dataset using the ConvNeXt-base backbone. The diagonal values represent the percentage of true positive predictions for each class.

### 3.3. Tracking

The resulting tracking performance is presented in Table 8. In total, 1,792 frames, corresponding to 110 minutes of recordings, were evaluated, containing 487 ground-truth (GT) insect tracks. The tracker generated 404 tracks (IDs), with 187 identity switches and 289 fragmentation events. Overall, a Multiple Object Tracking Accuracy (MOTA) of 76.1% with an Identity Precision of 79% (IDP), Identity Recall of 70% (IDR) and IDF1 of 74.4% was achieved. Most identity switches (IDSW = 106) and fragmentation events (FM = 163) occurred in recordings from camera Pi12, particularly during periods when three to six small bees were simultaneously present in the field of view. In contrast, the **RT-Impact** recordings, sampled at three-second intervals, achieved consistently high MOTA values of 84–90% across 64 GT tracks. These recordings primarily contained honeybees and bumblebees, which were captured at a higher image resolution than the bees in the **Orchard** recordings.

**Table 8:** Tracking evaluation showing the project and camera (CAM), number of frames (FR), ground-truth (GT) detections (det.) and tracks (trac.) for each recording sequence, together with the number of false-negative (FN) and false-positive (FP) detections, identity switches (IDSW), and fragmentation events (FM). Multiple Object Tracking Accuracy (MOTA), Identity Precision (IDP), Identity Recall (IDR) and IDF1 are also reported. The total values are calculated across all recording sequences based on the total number of GT detections, tracks, FN, FP, IDSW and tracked IDs.

| Project | CAM | FR | GT<br>det. | GT<br>trac. | IDs | FN | FP | ID-SW | FM | MOTA<br>(%) | IDP<br>(%) | IDR<br>(%) | IDF1<br>(%) |
| --- | --- | --- | --- | --- | --- | --- | --- | --- | --- | --- | --- | --- | --- |
| **RT-Impact** | Rt1a | 440 | 450 | 45 | 46 | 64 | 4 | 1 | 2 | 84.7 | 98.7 | 85.6 | 91.7 |
| **RT-Impact** | Rt1b | 100 | 123 | 18 | 18 | 4 | 0 | 9 | 2 | 89.4 | 94.1 | 91.1 | 92.6 |
| **Orchard** | Pi10 | 143 | 169 | 60 | 45 | 12 | 0 | 28 | 13 | 76.3 | 63.7 | 59.2 | 61.3 |
| **Orchard** | Pi11 | 181 | 169 | 31 | 29 | 25 | 57 | 7 | 9 | 47.3 | 66.1 | 78.7 | 71.9 |
| **Orchard** | Pi12 | 357 | 937 | 164 | 141 | 148 | 2 | 106 | 163 | 72.7 | 73.6 | 62.1 | 67.4 |
| **Orchard** | Pi13 | 113 | 109 | 4 | 8 | 34 | 4 | 9 | 6 | 56.9 | 53.2 | 38.5 | 44.7 |
| **Orchard** | Pi16 | 233 | 361 | 149 | 104 | 45 | 0 | 20 | 86 | 82.0 | 77.5 | 67.9 | 72.4 |
| **Orchard** | Pi17 | 225 | 227 | 16 | 13 | 23 | 0 | 7 | 8 | 86.8 | 89.7 | 80.6 | 84.9 |
| Total | All | 1792 | 2545 | 487 | 404 | 355 | 67 | 187 | 289 | 76.1 | 79.0 | 70.0 | 74.2 |

### 3.4. Computational speed and edge processing

The computational performance of the proposed pipeline was evaluated on several computing platforms, including edge-processing devices. These platforms included a standard workstation equipped with an Intel Xeon E5-2620 v4 @ 2.10 GHz CPU and an NVIDIA Quadro RTX 8000 GPU, a laptop-class system with an AMD Ryzen 7 PRO 6850U CPU (without GPU acceleration), and two edge devices: a Raspberry Pi 4 (RP4) with 4 GB RAM and a Raspberry Pi 5 (RP5) with 8 GB RAM.

Edge devices were selected due to their comparable cost and relevance for field deployment. The pipeline was evaluated by processing 35 images recorded with insect activity. Detailed runtime metrics are reported in Table 9. During testing, an external SSD containing the images was connected to the Raspberry Pi devices, which were accessed remotely over Wi-Fi without a connected display. The YOLO11s model was optimized using NCNN, a high-performance neural network inference framework, which resulted in improved performance only on the RP5 device.

**Table 9:** Processing time performance for the *InsectDCT* algorithm executed on a workstation with GPU, a CPU-based laptop, and Raspberry Pi (RP) devices. All values are reported as the average processing time per image (in seconds), computed over a test set of 35 images containing 39 insect crops. The total pipeline processing time using YOLO11s with EfficientNetV2-S when processing an image containing a single insect is also listed.

| Platform | YOLO11s | YOLO11s<br>NCNN | YOLO11m | ResNet50V2 | Efficient-<br>NetV2-S | ConvNeXt-<br>Base | Total time<br>YOLO11s+<br>EfficientNet |
| --- | --- | --- | --- | --- | --- | --- | --- |
| GPU | 0.047 | NA | 0.083 | 0.0079 | 0.0175 | 0.0171 | 0.088 |
| CPU | 0.614 | 0.627 | 1.51 | 0.036 | 0.0614 | 0.0913 | 0.729 |
| RP5 | 3.25 | 1.88 | 7.99 | 0.127 | 0.149 | 0.314 | 2.14 |
| RP4 | 9.79 | 9.91 | 25.5 | 0.310 | 0.400 | 0.883 | 10.3 |

### 3.5. Evaluation of pipeline with classification and tracking

Six days of recordings from different locations within the **Orchard** project were used to evaluate the entire pipeline, including tracking. The recordings were captured at a frame rate of 0.5 fps in HD resolution using Raspberry Pi 3 cameras. Each track was manually verified by inspecting the images contained, as illustrated in Figure 8. The tracks were manually categorized as correctly classified, containing multiple species, false positives, or unsure due to small or blurry insects that were too difficult to reliably classify. Tracks containing multiple species were treated as biologically valid for the main accuracy analysis, as the target taxon was still present within the majority of the crops in the track.

**Figure 8:**
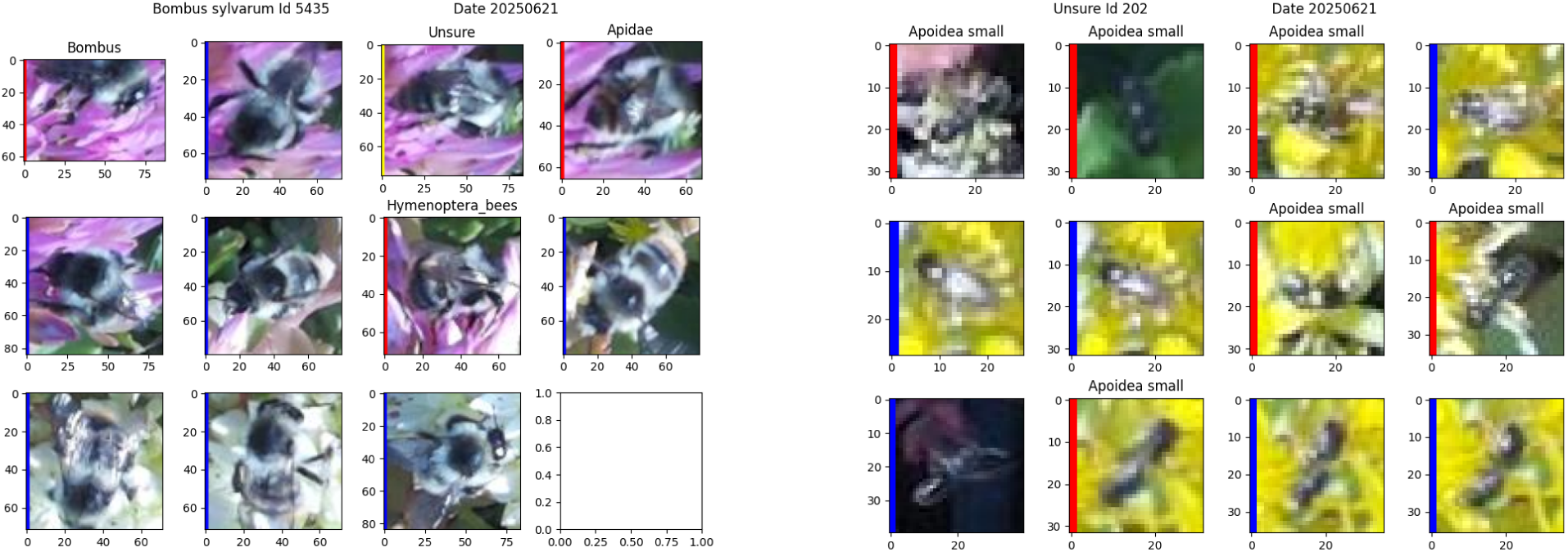
Shows examples of tracks with individual crops of detected and classified insects across image sequences. Left image crops contained in track of *Bombus sylvarum*. Right image crops contained in an “unsure” track with many crops classified as Apoidea small.

In total, 4,753 tracks were inspected, of which 3,469 were correctly classified. An additional 141 tracks contained multiple species, resulting in a total of 3,610 valid tracks and an overall precision of 75.95%. False-positive and unsure tracks accounted for 363 and 780 tracks, respectively. If tracks categorized as unsure are also accepted as correctly classified, the precision increases to 90.86%. The final distribution of validated tracks for a subset of images in the **Orchard** dataset shown in Figure 9 indicates that small insects, particularly the Apoidea small and hymenoptera groups, account for a large proportion of unsure tracks. This is mainly due to the very small crop sizes, often around 30 × 30 pixels, as illustrated in Figure 8.

**Figure 9:**
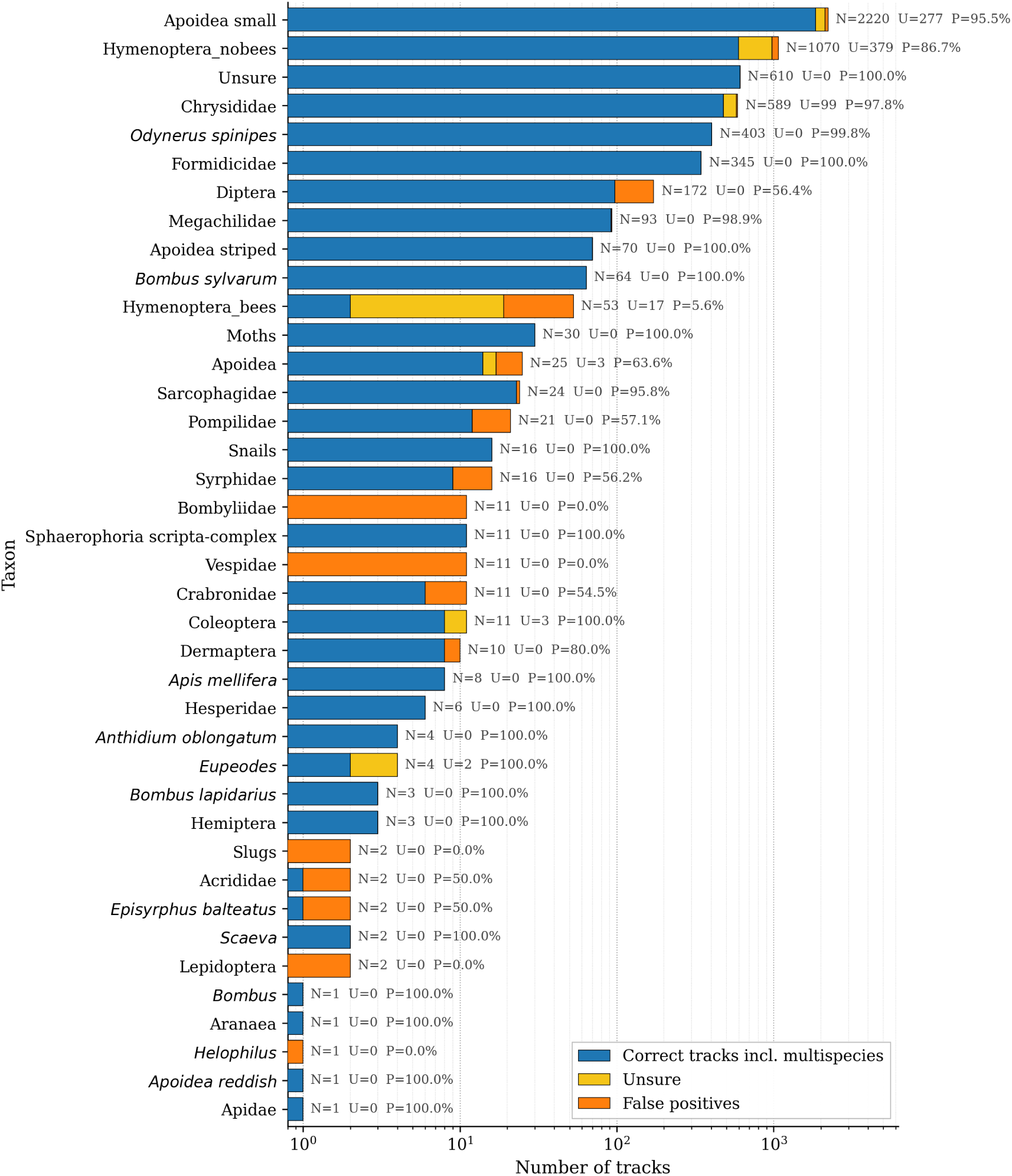
The number of tracks of insect taxa identified in recordings from six different camera locations from the *Orchard* dataset, each monitored over a one-day period. The correct tracks incl. multispecies are colored blue. Unsure tracks are shown in yellow and false positive tracks are shown in orange.

### 3.6. Monitoring insect abundance based on activity in camera recordings

The camera-derived activity data captured in the **Orchard** recordings show clear temporal patterns on both diel and seasonal scales. The daily activity curves differed between the pollinator groups (Figure 10): bees and butterflies showed predicted activity maxima around midday, peaking at approximately 13:20 h and 12:50 h, respectively, while hoverflies peaked earlier in the day at approximately 8:10 h. The estimated activity window above 50% of maximum activity was the broadest for hoverflies (12:20 h), followed by butterflies (9:00 h) and bees (7:10 h). Seasonal occurrence patterns also showed that the tracking data can resolve taxon-specific phenological windows throughout the sampling season (Figure 11).

**Figure 10:**
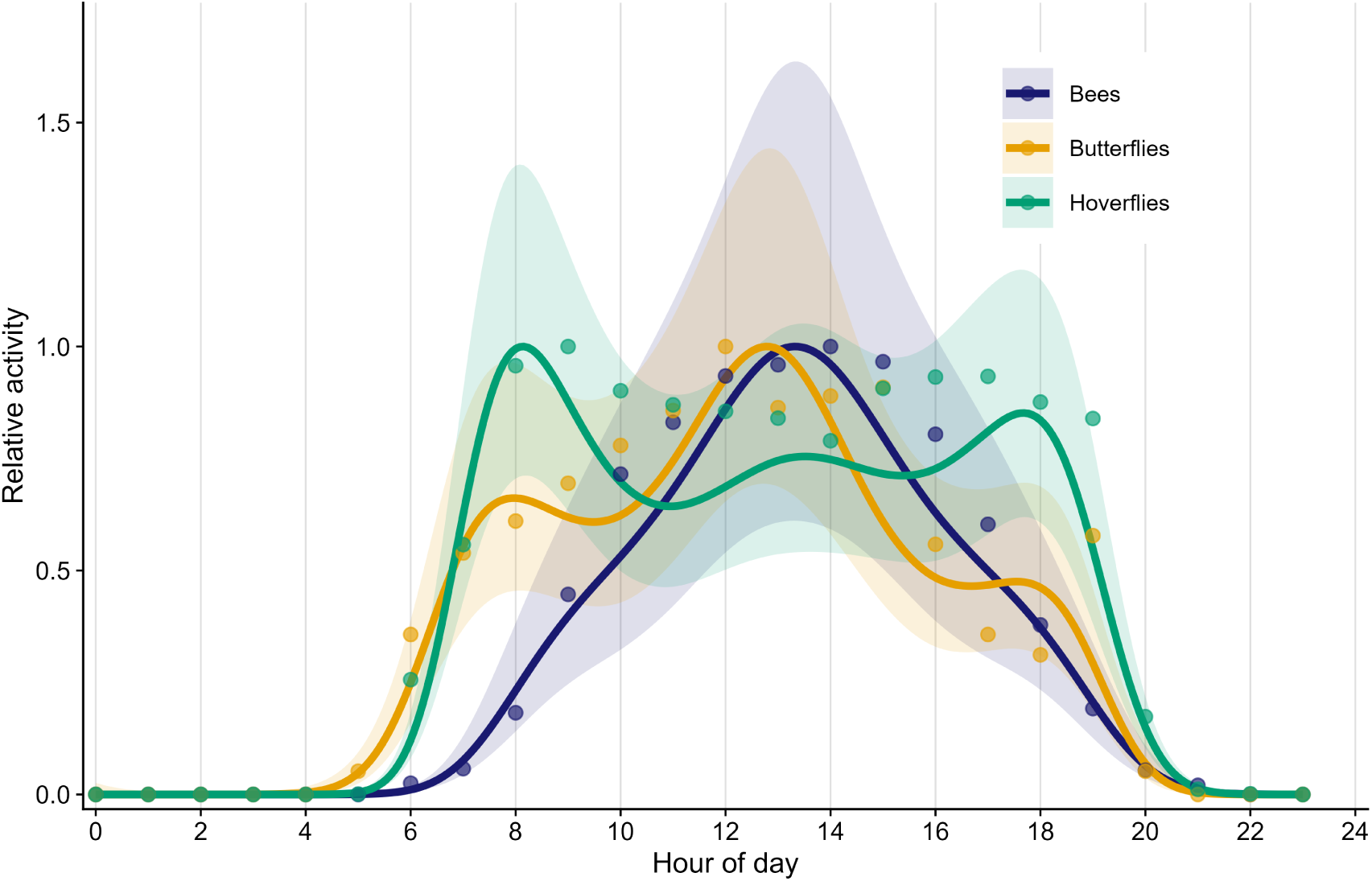
Daily activity patterns of butterflies, hoverflies, and bees in the all of the **Orchard** recordings. Smoothed curves show group-specific activity patterns across the day based on observation time. Points indicate hourly observation counts scaled to the maximum within each pollinator group. Shaded bands represent approximate 95% confidence intervals from negative-binomial GAMs. Activity is shown as relative activity to facilitate comparison of temporal patterns among groups rather than absolute abundance.

**Figure 11:**
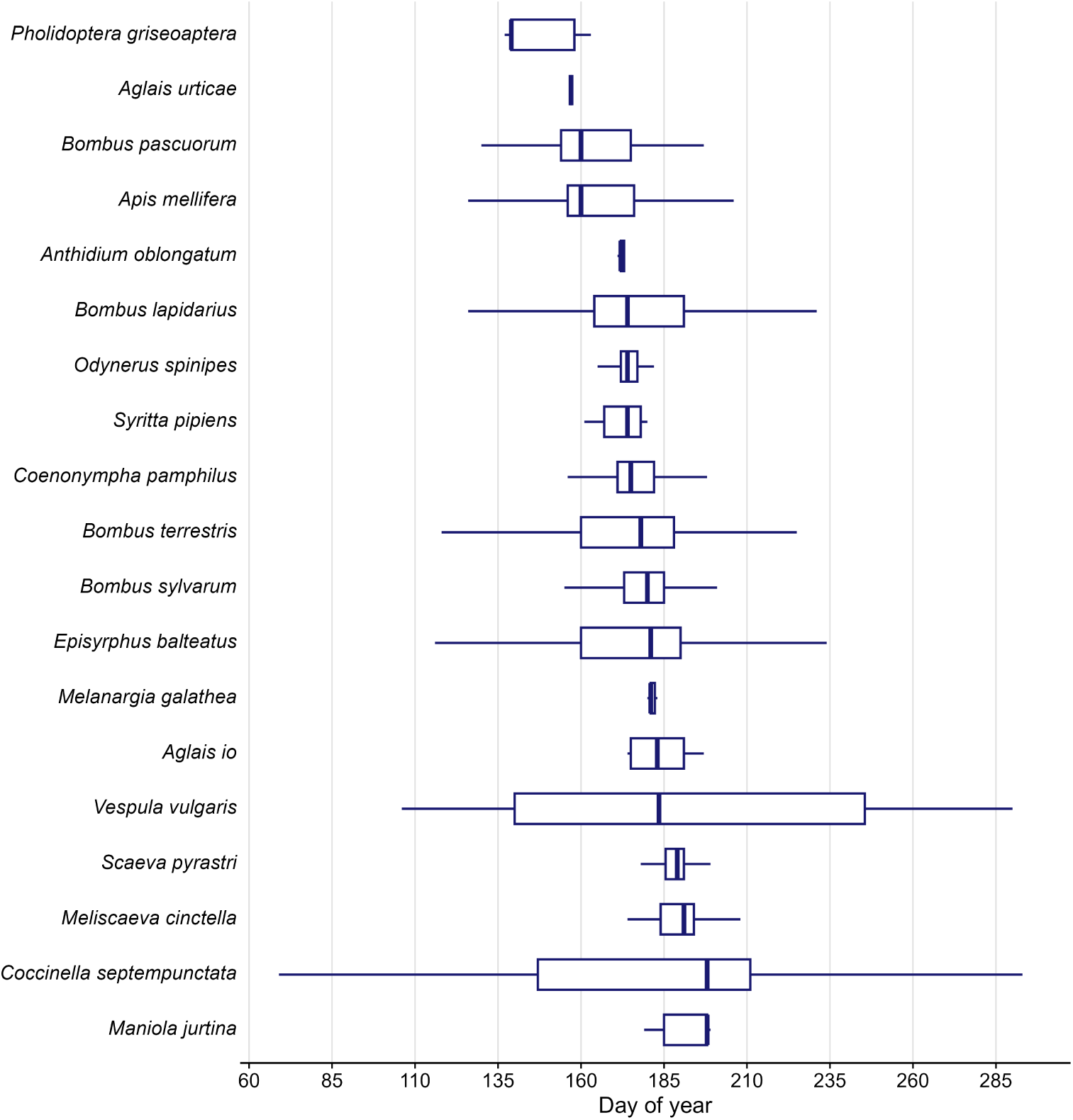
Phenology of observed insect species across the sampling season in the **Orchard** recordings. Horizontal boxplots show the distribution of observations for each species across day of year. Boxes indicate the interquartile range, whiskers show the observed temporal range excluding outliers according to the standard boxplot definition. Only taxa assigned to species level and represented by at least 20 observations are shown. Species are ordered by median day of observation.

## 4. Discussion and conclusion

This study presents *InsectDCT* as a generalized deep learning pipeline for detecting, hierarchically classifying, and tracking insects and other small animals in camera-trap recordings from natural and semi-natural floral environments. The pipeline addresses three central limitations of current automated insect-monitoring systems: limited transferability across field conditions, restricted taxonomic resolution, and reduced sensitivity to small or distant insects in complex natural scenes (Høye et al., 2025; Bjerge et al., 2024, 2023c; Svenning et al., 2026; Bjerge et al., 2023b). By combining a detector trained on many different vegetation communities and camera systems, a hierarchy-aware classifier covering 80 taxonomic and functional groups, and track-by-detection algorithm for temporally linked observations, *InsectDCT* provides a reusable framework for converting large image streams into standardized ecological observations.

A central contribution of the pipeline is its broad applicability across diverse scenes and recording conditions. Rather than being trained for a single plant species, camera setup, or local insect community, *InsectDCT* was developed using image data from multiple projects, camera systems, flowering plants, and vegetation backgrounds. Our results show that this diversity translates into robust performance on independent test datasets including unseen recording contexts and vegetation backgrounds. Another major contribution of this work is the development and release of two complementary datasets: one for insect detection and localization using both RGB and motion-enhanced images, and one for hierarchical taxonomic classification across multiple arthropod groups. These datasets are broader in background structure, camera type, and taxonomic composition than many previous camera-trap studies, which were often constrained to specific plant species, camera setups, or target taxa (Bjerge et al., 2023a, 2024, 2023c,b).

Beyond technical evaluation, field application demonstrates how *InsectDCT* can translate raw image streams into ecologically interpretable time series. The diel activity curves and phenological distributions extracted from the tracked detections are particularly relevant because pollinator activity varies throughout the day and season in response to taxon-specific thermal constraints, solar radiation, wind, and availability of floral resources (Corbet et al., 1993; Gilbert, 1985; Zoller et al., 2020; Mahon and Hodge, 2022; Gillespie et al., 2025). In this sense, the pipeline provides a non-lethal approach for quantifying relative activity, flower visitation, and phenology of common flower-visiting taxa at temporal resolutions that are difficult to achieve with repeated manual observations (Naqvi et al., 2022; Ratnayake et al., 2021; Høye et al., 2025).

However, it should be taken into account that camera-derived counts should be interpreted as activity or visitation indices rather than direct estimates of population size unless additional assumptions are met (Høye et al., 2025; Rowcliffe et al., 2008, 2014; MacKenzie et al., 2002). Therefore, *InsectDCT* outputs are best treated as standardized, camera-specific activity-density or visitation-rate proxies unless they are calibrated against independent counts or analyzed with models that explicitly account for detection probability and sampling effort.

Svenning et al. (2026) proposed a generic method for arthropod detection trained on a large dataset, allowing for scale and size-agnostic detection; however, the approach was developed using relatively homogeneous backgrounds. Previous studies (Bjerge et al., 2023a, 2024) have demonstrated insect detection in natural environments, but were limited to models trained primarily on the backgrounds of *Sedum* plants. The detector presented in this work builds on the approach in (Bjerge et al., 2023b), extending it with a substantially greater diversity of vegetation backgrounds. This increased variability improves the robustness of the model and reduces the likelihood of false-positive detections in complex natural scenes. To our knowledge, *InsectDCT* is therefore among the most broadly trained and tested camera-based insect monitoring pipelines for natural vegetation backgrounds currently available, which makes it especially valuable for users who want to apply automated insect monitoring in new projects without first developing a fully project-specific detection algorithm from scratch.

The detector trained on motion-enhanced images achieved a higher average F1-score than the RGB-based detector and increased average recall, while also showing lower variation in F1-score among projects. The strongest improvements occurred in particularly challenging backgrounds, such as the **GreenHouse**, **MAMBO**, and **PollWatch** datasets, where recall increased substantially when motion-enhanced images were used. These results support the findings of Bjerge et al. (2023b), demonstrating that models trained with motion-enhanced images generalize better to new environments and domains.

Designing an appropriate camera setup for genus- or species-level insect identification involves a trade-off between frame rate, camera resolution, field of view, and the resulting size of insect crops. A larger field of view increases the likelihood of capturing insects within the scene and allows for monitoring larger floral patches, but it also reduces the pixel size of individual insects and can therefore lower classification reliability unless higher-resolution cameras are used. In contrast, close-range imaging can improve detection and classification by producing larger insect crops, but at the cost of a smaller sampling area. This trade-off is illustrated by previous camera-based monitoring systems. Varga-Szilay et al. (2024) detected and tracked bumblebees visiting flowers, but the relatively large distance between cameras and flowers made it difficult to obtain a reliable classification for diverse insect communities. Smith et al. (2026) presented AutoPollS, which uses four Borescope USB cameras placed close to the plants and only accepted detections larger than 120 × 120 pixels. This design increases the robustness of detection and classification, but the resulting field of view is considerably narrower than in the datasets presented here.

In contrast, the detector in *InsectDCT* was trained specifically for small insects, with an average object size of 71 × 66 pixels (standard deviation 36 × 33) in HD-resolution images. Robust detection at this scale allows *InsectDCT* maintain a broader field of view while being able to detect and classify smaller pollinators that would otherwise be missed or excluded. At the same time, this specialization introduces a scale-related trade-off: performance decreases for larger insects, which were less well represented in the small-object-focused training data. To address this, a separate version of the color-based detector was trained on additional images that contain larger specimens. However, this model performed less effectively on small insects, highlighting the difficulty of developing a single detector that performs equally well across the full range of insect body sizes and image scales, even though YOLO architectures are designed for multi-scale object detection.

A key strength of the *InsectDCT* classifier is its hierarchical structure, which makes the output useful even when fine-grained identification is uncertain. Instead of forcing every detection into a species-level label, the model can retain ecologically informative predictions at higher taxonomic levels, such as order or family, while providing genus- or species-level classifications where the image quality and taxonomic separability allow it. The higher image quality and fewer insects may have contributed to the improved tracking performance observed in the RT-Impact recordings. In the current implementation, the classifier already covers a broad range of arthropod groups, including the main flower-visiting taxa observed in the recordings, with bees, hoverflies, and butterflies represented at a relatively fine taxonomic resolution. Nevertheless, the dataset used to train the hierarchical classifier exhibits a long-tailed class imbalance. To compensate for this imbalance during training, a balanced softmax loss function was used. However, the method only partially mitigates the imbalance and classes with very few training samples remain challenging for the model to classify accurately. The hierarchical classifier could be further refined by resolving broad groups such as Coleoptera and Hemiptera into lower taxonomic ranks. Currently, these categories contain a high diversity of visually distinct taxa, which reduces the classification accuracy.

We recommend extending the dataset with additional taxonomic groups when applying the pipeline to environments containing insect species not represented in the current recordings. Even if species-level predictions are likely unreliable for previously unseen taxa or regions, the pipeline can still assign detections to broader taxonomic groups and generate standardized image crops that are already partially sorted. These outputs can then be reviewed and re-annotated more efficiently, providing a structured starting point for extending the classifier with additional taxa rather than requiring researchers to build a training dataset from unfiltered raw images. In this sense, *InsectDCT* can support both immediate ecological inference at coarser taxonomic levels and the iterative development of more region- or project-specific classifiers (Bjerge et al., 2023c).

Tracking is a central part of *InsectDCT* since it converts frame-level detections into standardized activity and visitation indices. By reducing the repeated counting of the same individual within short image sequences and generating track-level summaries such as visit duration, arrival time, and predominant taxonomic identity, the pipeline provides outputs that are directly relevant for estimating relative activity, visitation rates, diel patterns, and pollinator behavior.

Our evaluation of the complete tracking pipeline was conducted on a highly challenging dataset with an average insect image size of only 50 × 50 pixels, making it difficult to reliably identify many small solitary bees. The tracking algorithm also struggles when small insects cross paths, which can result in mixed or fragmented tracks. However, for video recordings, we have also experimented with YOLO11 ByteTrack (Zhang et al., 2022) or BoT-SORT (Aharon et al., 2022) tracking algorithms in the initial processing stage. In this approach, insect detection and tracking are combined in a single step, followed subsequently by taxonomic classification. However, preliminary testing on eight video sequences indicated inferior performance compared to our approach (results not shown). In particular, occlusions and missed detections often caused a single insect to be represented by multiple fragmented tracks, reducing overall tracking accuracy.

For custom insect camera systems based on Raspberry Pi 4 or 5 platforms (Sittinger et al., 2024; Ratnayake et al., 2021; Preti et al., 2021), our pipeline enables near-real-time processing of time-lapse recordings. On the Raspberry Pi 5, detection requires approximately 1.88 s per image, increasing to 2.14 s when classification is included (approximately 0.47 fps), with tracking adding relatively little computational overhead. This processing rate exceeds the 0.33 fps at which real-time insect tracking has previously been demonstrated (Bjerge et al., 2023a), suggesting that the complete pipeline can operate directly on a Raspberry Pi 5, although with limited computational headroom. Such on-device processing could enable selective storage of images containing detected insects, reducing data storage requirements. In contrast, the Raspberry Pi 4 does not achieve the processing speed required to maintain a sampling rate suitable for reliable tracking under the tested conditions. For deployments with lower temporal resolution, tracking may instead be omitted, as time-lapse sampling has been shown to provide a viable alternative to track-based monitoring when sufficiently long sampling intervals are used (Bjerge et al., 2025). Additionally, reducing the detection confidence threshold can increase recall to over 90% of insects present in the recorded footage, although potentially at the expense of precision (Table 6).

As an alternative to deploying the complete pipeline on a camera trap equipped with a Raspberry Pi, and to address the computational constraints of edge deployment, we recommend a two-stage processing strategy as proposed by Smith et al. (2026). In the first stage, insect detection using the YOLO11s NCNN model is performed in real time on the camera device. This can be achieved on a Raspberry Pi 5 for time-lapse intervals as short as 1.88 seconds, or on a Raspberry Pi 4 for intervals of 10–60 seconds. Only images containing detected insects are stored locally and subsequently processed offline using the full *InsectDCT* pipeline for taxonomic classification and tracking. This approach exploits the relatively low computational cost of optimized YOLO-based detection on edge hardware while offloading the classification and tracking tasks to systems with greater flexibility and processing capacity. Consequently, power consumption, storage requirements, and data transmission needs in the field can be substantially reduced without compromising taxonomic resolution or tracking performance.

Users wishing to deploy *InsectDCT* in a new ecological context, should follow a structured workflow that spans image acquisition, annotation, model training, and validation. In the field acquisition stage, cameras should be positioned at a distance of 30 cm with a wide-angle lens and 40-50 cm with a 35mm focal lens (Pi HQ Camera) of the floral patch and any angle if the insect is within a detectable size range; based on the performance analysis presented in this study, a minimum crop size of approximately 50 × 50 pixels in HD resolution is advisable for reliable classification. Time-lapse intervals of 60 seconds or less are required for the motion-enhanced image pipeline. For video-based deployments, frame rates above 0.33 fps are required to maintain high performance of the tracking of individuals.

Images should be collected throughout the entire phenological season, under varying illumination conditions and at multiple locations, to ensure that the resulting data set reflects the natural variability of the target environment. For detection, the existing trained YOLO11 model can be applied directly in a first pass; images flagged as containing insects should be manually verified to remove false positives and recover missed detections. This reviewed set should ideally be incrementally added to the training corpus for iterative re-training following the procedure described in Section “Dataset for insect detection”.

For taxonomic classification, new taxa can be incorporated by supplementing the data described in Tables 3 and 4 with additional labels L1, L2 and L3, provided that a minimum of approximately 100–300 annotated crops per class are available; classes with fewer samples can be supplemented with images recorded with smartphones (e.g., GBIF images), keeping in mind the domain gap between such images and camera-trap crops discussed above. Finally, tracking parameters, particularly the cost threshold and the minimum number of detections per valid track, may require adjustment depending on the frame rate and typical insect movement patterns in the new recording setup. *InsectDCT* architecture, in which detection, classification, and tracking operate as independent stages with standardized inputs and outputs, is specifically designed to facilitate these adaptations, allowing researchers to replace or retrain individual components without modifying the rest of the pipeline.

The pipeline has the potential to support the extraction of ecological and behavioral foraging data for insects, allowing studies of e.g. insect flower visitation rates, flower handling times, and movement patterns among inflorescences that would be difficult or labor-intensive to obtain through manual observation alone. By linking consecutive detections into tracks, *InsectDCT* provides a temporal dimension to camera-trap data that goes beyond simple presence/absence or abundance estimates. This information is relevant for studies of plant-pollinator interaction networks, where species identity, visitation frequency, and movement patterns could be used to automatically infer the functional contribution of each pollinator species.

Despite its strong performance under diverse field conditions, *InsectDCT* has several limitations that should be acknowledged. The taxonomic classifier is currently trained on Northern and Central European pollinator taxa, with a large part of the dataset focused on vegetation dominated by plant species of the *Sedum* genus, which implies that the pipeline will require re-training for direct transferability to other geographic regions or habitat and vegetation types. Researchers deploying the pipeline in new contexts are encouraged to supplement the existing dataset with locally collected and annotated images, following the incremental re-training workflow described above, which has been specifically designed to accommodate this kind of extension. The classes with few training samples, such as rare solitary bees and some folded-wing Lepidoptera, show reduced classification accuracy, and this imbalance could be mitigated in future versions by re-training the algorithm by users who require improved performance in those insect groups. Regarding the tracking algorithm, it can produce fragmented or merged trajectories when small insects cross paths or are temporarily occluded, and tracks should therefore be interpreted as activity proxies rather than records of unique individuals over extended periods.

Taken together, these limitations do not reduce the broader utility of this pipeline, but instead define clear pathways for extension. Because *InsectDCT* combines open datasets, modular detection and classification models, anomaly handling, and track-level output, it can serve both as an immediately usable monitoring tool and as a foundation for project-specific adaptation. By lowering the amount of annotation and algorithm development required before camera-based insect monitoring can be applied in new ecological contexts, *InsectDCT* can help make high-resolution, non-invasive pollinator monitoring more scalable, comparable, and accessible across studies.

## CRediT authorship contribution statement

Kim Bjerge: Conceptualization, Investigation, Data curation, Project administration, Methodology, Software, Validation, Visualization, Writing – original draft. Simon F. A. Wogram: Conceptualization, Investigation, Data curation, Methodology, Software, Validation, Visualization, Writing – review & editing. Pau Enric Serra-Marin: Investigation, Data curation, Writing – review & editing. Otar Sakhiashvili: Writing – review & editing. Andreas Bonnerup Jepsen: Data curation, Validation. Toke T. Høye: Funding acquisition, Writing – review & editing.

## Acknowledgments

This research was partly funded by the European Union’s Horizon Europe Research and Innovation programme, under Grant Agreement No. 101060639 (**MAMBO**). The authors would like to thank all participants in the **MAMBO** project for their valuable contributions to the collection of image recordings throughout Europe. We also thank Hjalte Ro Poulsen from the Department of Geosciences and Natural Resource Management, University of Copenhagen, for providing images from the **Heather** project.

## Data availability

Training and testing results presented in this paper are based on version 6 of the detector and classifier datasets (DV6 and CV6). These datasets and the GT dataset for tracker performance evaluation can be downloaded from:

https://zenodo.org/records/21154490

The source code and trained models for the deep learning pipeline can be downloaded from:

https://github.com/kimbjerge/insectDCT

## Declaration of Generative AI and AI-assisted technologies in the writing process

During the preparation of this work, the author(s) used ChatGPT in order to improve formulations in writing. After using this tool/service, the authors reviewed and edited the content as needed and take full responsibility for the content of the publication.

## Supporting information

Figures S1–S6 present example images from the detection dataset, illustrating the diversity of plant backgrounds in several projects to train a robust and generalized insect detector.

**Fig. S1:**
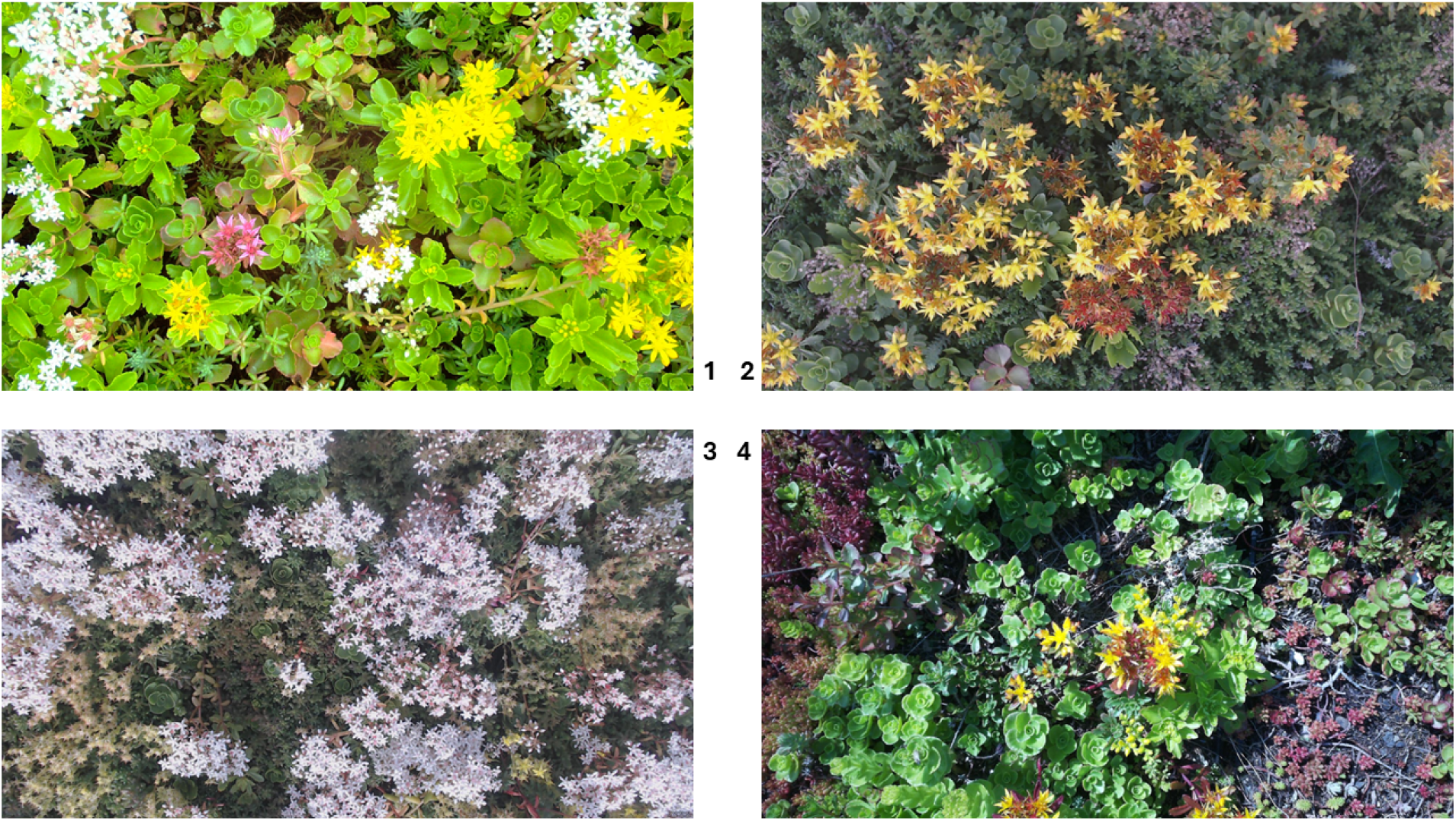
Examples of *Sedum* plants are presented from the **GreenRoof** project (1–3) and the **Orchard** project (4).

**Fig. S2:**
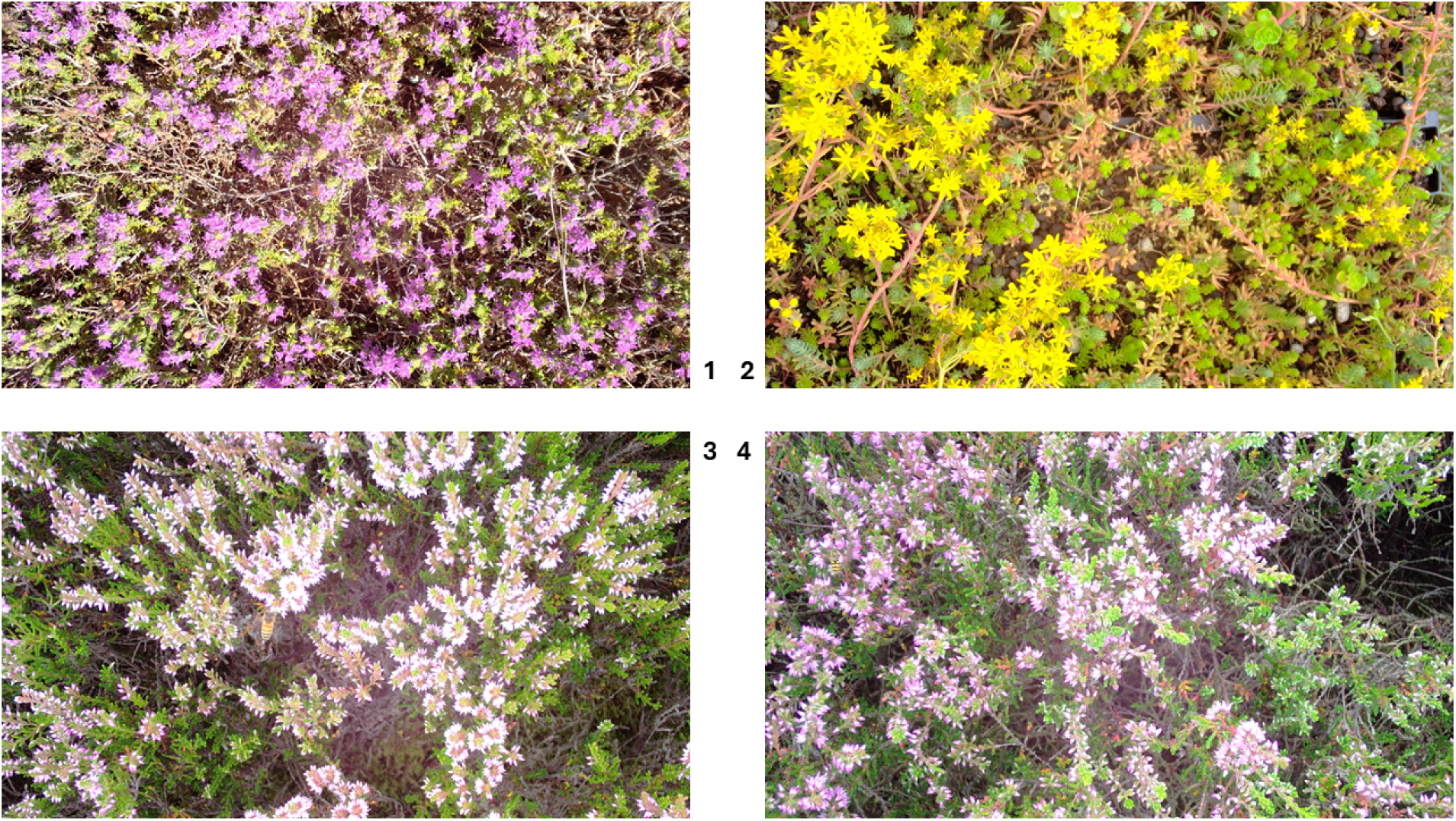
Examples of Herbs, *Sedum* and Heather plants are presented from the **MAMBO** project (1,2) and the **Heather** project (3,4).

**Fig. S3:**
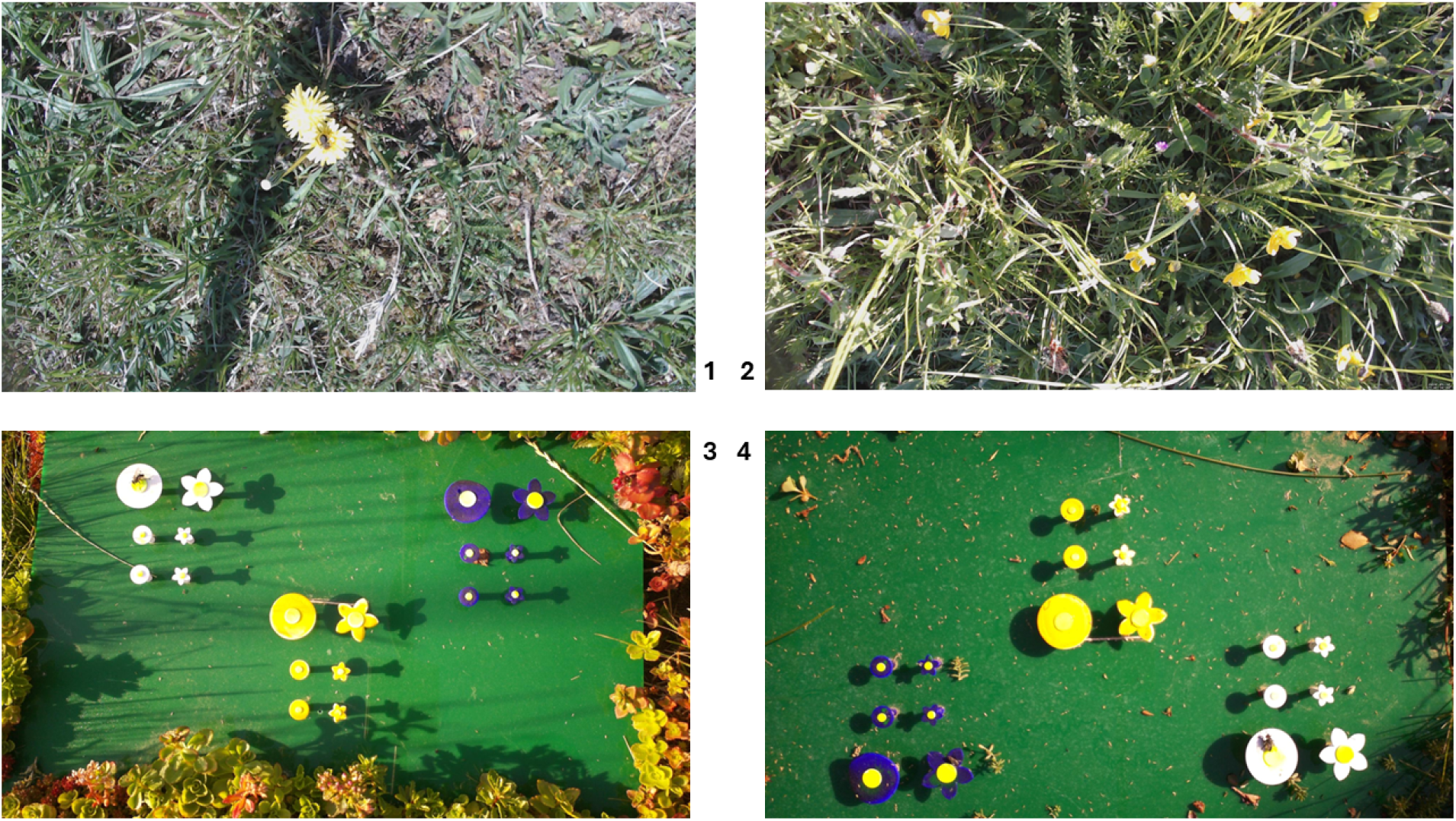
Examples of mixed plants and artificial flowers are presented from the **PollWatch** (1,2) and **Orchard** (3,4) projects.

**Fig. S4:**
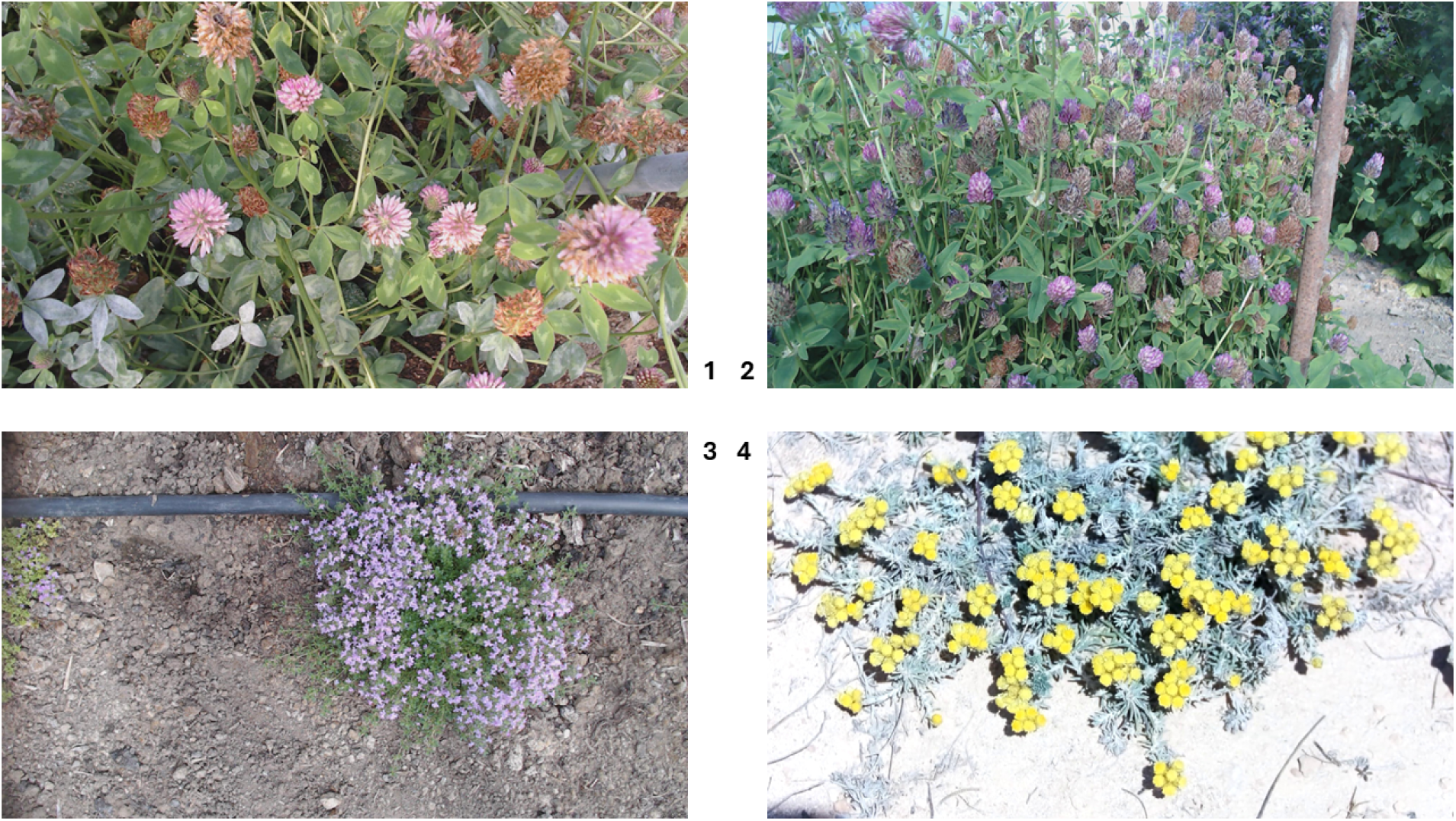
Examples of Red Clover, Sea Rocket plants are presented from the **GreenHouse** (1-3) and **ACSHQ** (4) projects.

**Fig. S5:**
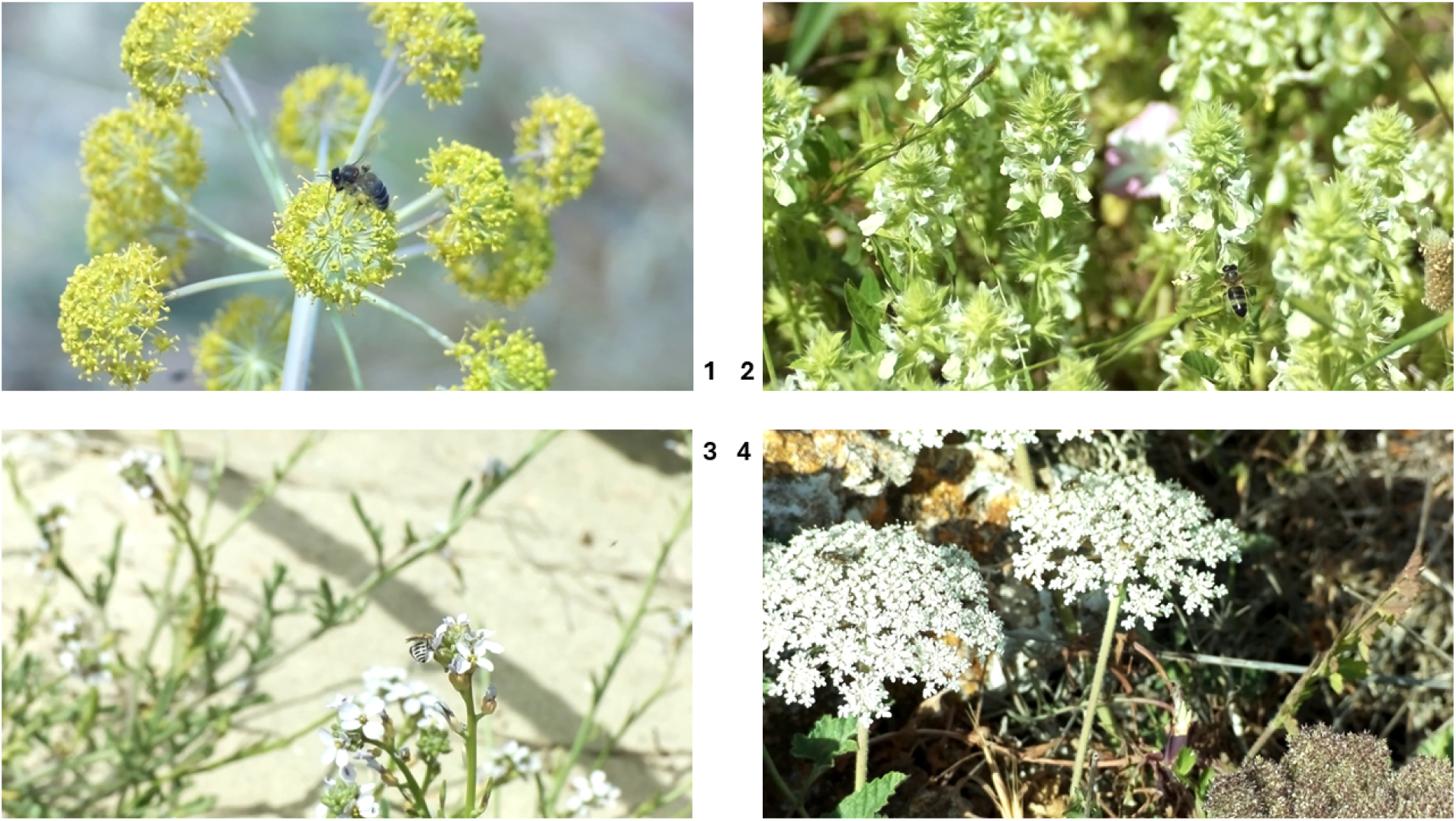
Examples of different plants with insects are presented from the **ACSHQ** project.

**Fig. S6:**
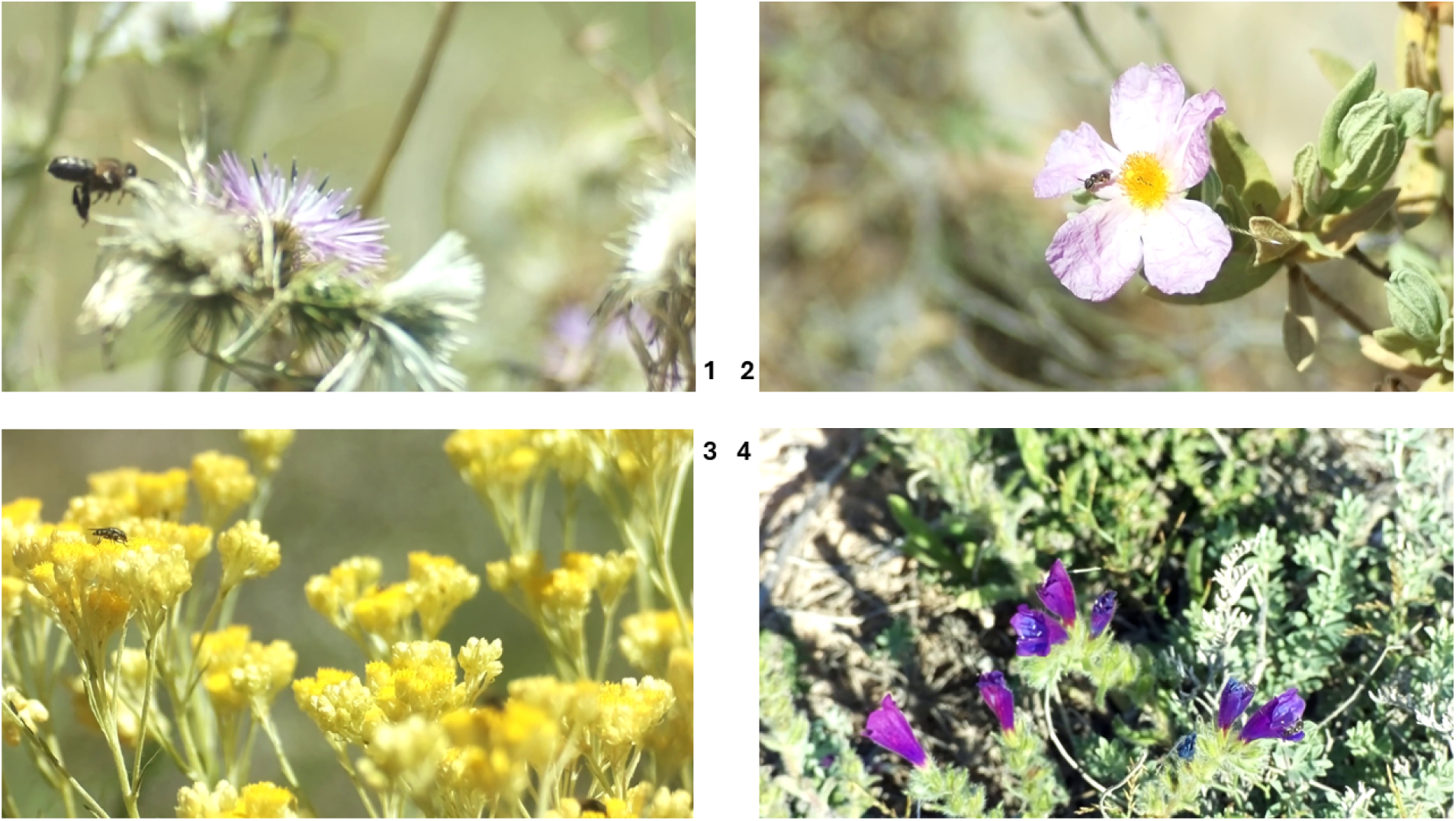
Examples of different plants with insects are presented from the **ACSHQ** project.

